# A TLR7 Agonist Conjugated to a Nanofibrous Peptide Hydrogel as a Potent Vaccine Adjuvant

**DOI:** 10.1101/2024.03.07.583938

**Authors:** Erin M. Euliano, Brett H. Pogostin, Anushka Agrawal, Marina H. Yu, Tsvetelina H. Baryakova, Tyler P. Graf, Jeffrey D. Hartgerink, Kevin J. McHugh

## Abstract

Toll-like receptors (TLRs) recognize pathogen- and damage-associated molecular patterns and, in turn, trigger the release of cytokines and other immunostimulatory molecules. As a result, TLR agonists are increasingly being investigated as vaccine adjuvants, though many of these agonists are small molecules that quickly diffuse away from the vaccination site, limiting their co-localization with antigens and, thus, their effect. Here, the small-molecule TLR7 agonist 1V209 is conjugated to a positively-charged multidomain peptide (MDP) hydrogel, K_2_, which was previously shown to act as an adjuvant promoting humoral immunity. Mixing the 1V209-conjugated K_2_ 50:50 with the unfunctionalized K_2_ produces hydrogels that retain the shear-thinning and self-healing physical properties of the original MDP, while improving the solubility of 1V209 more than 200-fold compared to the unconjugated molecule. When co-delivered with ovalbumin as a model antigen, 1V209-functionalized K_2_ produces antigen-specific IgG titers that were statistically similar to alum, the gold standard adjuvant, and a significantly lower ratio of Th2-associated IgG1 to Th1-associated IgG2a than alum, suggesting a more balanced Th1 and Th2 response. Together, these results suggest that K_2_ MDP hydrogels functionalized with 1V209 are a promising adjuvant for vaccines against infectious diseases, especially those benefiting from a combined Th1 and Th2 immune response.

**Table of Contents:** Activation of toll-like receptors (TLRs) stimulates a signaling cascade to induce an immune response. A TLR7 agonist was conjugated to an injectable peptide hydrogel, which was then used to deliver a model vaccine antigen. This platform produced antibody titers similar to the gold standard adjuvant alum and demonstrated an improved balance between Th1- and Th2-mediated immunity over alum.

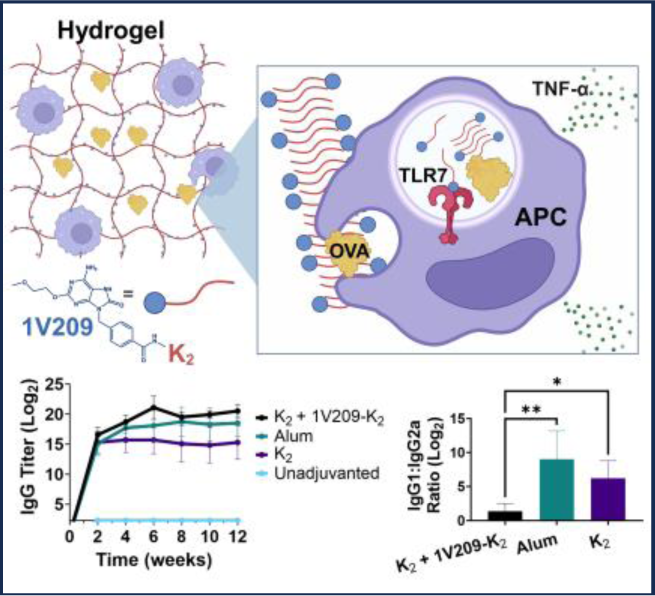

## Introduction

Vaccines are, on the whole, immensely effective at preventing illness and mortality, yet potent vaccines against some of the most widespread pathogens remain elusive. One promising approach to overcoming the challenges faced in the development of vaccines against existing and emerging pathogens is the implementation of novel immune adjuvants. Adjuvants can enhance vaccine efficacy by increasing antigen immunogenicity, controlling antigen delivery, and/or directing the phenotype of the effector response.^[1]^ Since the first adjuvant, alum (aluminum hydroxide microparticles), was developed in the 1920s, only eight additional adjuvants have been included in FDA-approved vaccines.^[2]^ Unfortunately, our current library of antigens and adjuvants has yet to be combined in a way that provides widespread sterilizing immunity for diseases such as tuberculosis, malaria, and human immunodeficiency virus (HIV) infections, highlighting the need for new and more effective vaccine adjuvants.

In order to develop vaccines that confer protective immunity, vaccine formulations must activate the appropriate immunological pathways. The two major arms of the adaptive immune response are cellular immunity (Th1), associated with antigen-specific cytotoxic T cells, and humoral immunity (Th2), associated with B cell maturation and antibody production. Depending on the type of pathogen, one or both responses may be necessary for protection.^[3,4]^ The type of immune response generated from a vaccine or pathogen is dependent, in part, on the activation of the innate immune system. Specific pattern recognition receptors (PRRs) in antigen-presenting cells (APCs) recognize pathogen-associated molecular patterns (PAMPs), which are foreign substances common to broad sets of pathogens. The most well-studied family of PRRs are toll-like receptors (TLRs), which recognize PAMPs, such as bacterial cell wall lipopolysaccharide (LPS), double-stranded RNA, and flagellin. When these PAMPs are recognized by TLRs, APCs produce cytokines that direct the adaptive immune response to the correct phenotype to effectively clear the infection.^[5]^

There has been growing interest in using TLRs to direct the adaptive immune response in adjuvant development. To date, there have been two TLR agonists included in FDA-approved vaccine formulations: CpG 1018, a TLR9 agonist, and monophosphoryl lipid A (MPL), a TLR4 agonist that is a synthetic analog of LPS.^[2]^ These adjuvants have been used in the clinic to generate robust Th1 and Th2 responses to pathogens, including hepatitis B.^[6–8]^ Unfortunately, these compounds have not been able to confer long-lived, protective immunity against all pathogens when co-administered with current antigens.^[9]^ Instead, the development of new adjuvants that leverage the eight other TLRs that have not yet been used in FDA-approved vaccines may hold promise for enhancing vaccine efficacy.^[10]^

TLR7, which is expressed in the endosome of APCs, natively recognizes single-stranded RNA but can also be potently activated with small-molecule agonists, making it a particularly attractive target for adjuvants.^[11]^ Small-molecule adjuvants are desirable for their facile large-scale synthesis compared to large complex biomolecules like MPL and can be readily incorporated into existing adjuvant and vaccine platforms.^[12]^ Although there are many small-molecule TLR7 agonists in clinical and preclinical development, their utility is limited by their poor solubility and short *in vivo* half-life.^[13,14]^ Several groups have developed strategies to overcome these challenges and improve their adjuvancy by covalently attaching small-molecule TLR7 agonists to antigens or other adjuvants. For instance, covalently conjugating the small-molecule TLR7 agonist 1V209 to carbohydrates has been shown to improve its activity and promote a robust humoral immune response against the model antigen ovalbumin (OVA) in mice.^[15,16]^ Structurally similar hydrophobic small-molecule TLR7/8 agonists have been directly conjugated to peptide antigens to form nanoparticles for improved cancer immunotherapy^[17,18]^ and formulated in a synthetic polymer supramolecular hydrogel for sustained co-delivery with an influenza antigen.^[19]^ These studies use the hydrophobicity of the small-molecule agonist to create amphiphilic structures that self-assemble into nanostructures, thereby improving the overall solubility, bioavailability, and half-life of the compound. The inclusion of small-molecule TLR7 agonists into immunomodulating materials is a promising strategy to overcome some of the issues conventionally associated with their delivery and achieve the desired adaptive immune response.

Biomaterials fabricated from self-assembling peptides have shown promise as immunomodulatory materials for vaccine applications. For example, seminal work by the Collier group has shown that the short self-assembling peptide Q11 (Ac-QQKFQFQFEQQ-Am), which forms β-sheet-rich peptide nanofibers, possesses strong immunomodulatory properties and can serve as an adjuvant for a wide variety of antigens.^[20–23]^ Both the sequence and higher-order structure of these peptides play a critical role in directing and strengthening the immune response to the co-delivered antigen of interest.^[24,25]^ Although this approach can be very effective in some cases, implementing it requires the existence of a short peptide that confers protection against a pathogen as well as *a priori* knowledge of that peptide’s sequence. It can also be synthetically challenging to implement, requiring the formation of covalent bonds between the vaccine antigen and the peptide adjuvant to prolong their release. Additionally, these peptide antigens are often less immunogenic and limit the number of available epitopes for the immune system to recognize. To address these challenges, self-assembling peptide hydrogels that can physically entrap native whole antigens to delay their release have been investigated as vaccine adjuvants. Recent studies have shown that peptides formulated as a hydrogel can elicit stronger immune responses than the same sequences in solution.^[26,27]^ These materials have also been used to enhance the immunogenicity of clinically relevant vaccines, such as a West Nile Virus subunit antigen.^[28]^ As the field of immunomodulatory self-assembling peptides has matured, there has been a growing interest in combining the immunostimulatory properties of TLR agonists with self-assembled peptide nanofibers to enhance the immune response to vaccine antigens. Nanofiber-forming peptides that act as TLR agonists and hydrogels loaded with TLR agonists have been previously used as vaccine adjuvants.^[29–31]^ These systems, however, have frequently been limited by their inability to extend both antigen and adjuvant delivery simultaneously, which is critical for improving vaccine-induced immune responses.^[19]^

Multidomain peptide (MDP) hydrogels are a class of self-assembling peptides that have shown promise as immunomodulatory materials.^[32]^ These peptides have an amphiphilic core of six serine-leucine (SL) repeats flanked by two charged amino acid residues on either terminus. The physical properties of MDP hydrogels are easily tuned by altering the peptide sequence in the amphiphilic core or, more commonly, in the charged domain.^[33–36]^ These viscoelastic materials are shear-thinning and self-healing, allowing them to behave like a liquid when passing through a small-bore (31-gauge) needle, yet rapidly form a hydrogel *in situ* after injection. Previous investigations have demonstrated that the positively charged peptide Ac-KKSLSLSLSLSLSLKK-Am (abbreviated as K_2_) strongly biases the adaptive immune response to a co-delivered antigen towards humoral immunity.^[37]^ The immune response to OVA co-delivered with K_2_ was found to be stronger than OVA co-delivered with MDPs composed of flanking Glu (E_2_) and Arg (R_2_) residues, suggesting that K_2_ is the most promising sequence for further investigation. In the current study, we aim to explore the influence of modifying the N-terminus of the K_2_ MDP hydrogel with the small-molecule TLR7 agonist 1V209. We hypothesized that conjugating 1V209 to the K_2_ hydrogel would enhance the small molecule’s solubility, extend its *in vivo* persistence at the site of injection, and synergize with the immunostimulatory properties of the MDP to elicit a robust immune response (Figure 1).

**Figure 1.**
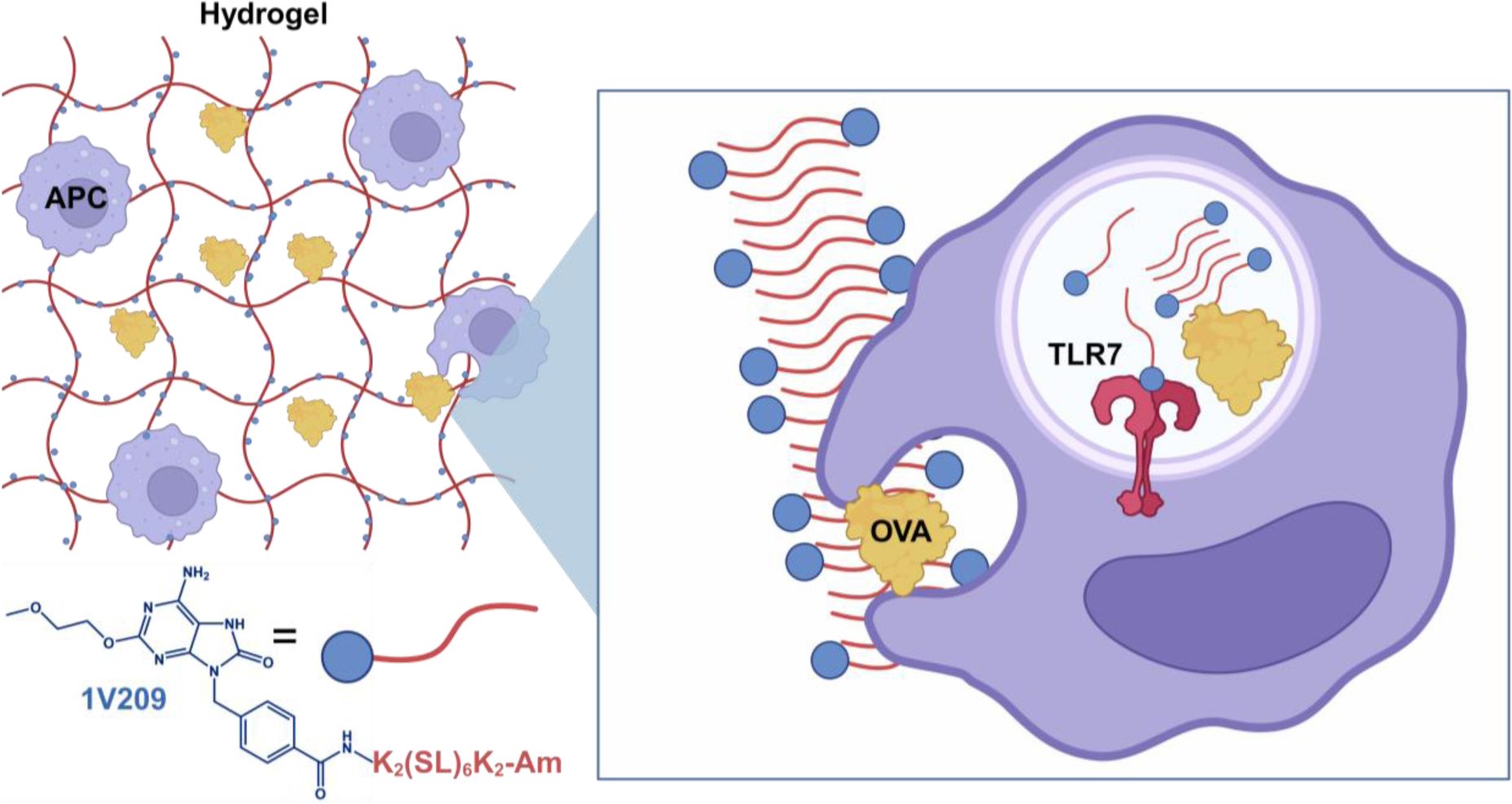
Schematic showing antigen-presenting cells (APCs) phagocytosing ovalbumin (OVA), a model antigen, with peptide nanofibers composed of a K_2_ self-assembling peptide functionalized with 1V209 on the N-terminus. After internalization, the 1V209 portion of the molecule interacts with TLR7 on the endosomal membrane and activates the innate immune system to enhance the adaptive immune response to OVA.

## Methods

### Solid-phase peptide synthesis (SPPS) and purification

Peptides were synthesized manually using standard Fmoc-based chemistry. Rink Amide MBHA resin and Fmoc-protected amino acids were supplied by Novabiochem (Millipore Sigma, Burlington, MA). Fmoc-protected low-loading MBHA rink-amide resin (1 equiv.) was deprotected with two washes of excess 25% v/v piperidine in a 1:1 mixture of dimethyl formamide (DMF) and dimethyl sulfoxide (DMSO) for 5 min each. Fmoc-protected amino acids (4 equiv.) were then preactivated for 1 min with hexafluorophosphate azabenzotriazole tetramethyl uronium (HATU; 3.95 eq.) and diisopropylethylamine (DIEA; 6 eq.) in a minimal amount of 1:1 DMF:DMSO then added to the resin for 20-30 min at ambient temperature. This process of deprotection and subsequent coupling was repeated for the addition of each subsequent amino acid residue until the peptide was complete. Several modifications were made to the base K_2_ peptide to yield three variants with a PEG spacer, two fluorescent variants, and the 1V209 variant. Coupling 5-(and 6)-Carboxytetramethylrhodamine (TAMRA) mixed isomers (Thermo Scientific, Waltham, MA) and Fmoc-8-amino-3,6-dioxaoctanoic acid (Fmoc-PEG2-OH) (Ambeed, Arlington Heights, IL) to peptides was performed using the same protocol as all other amino acids. To synthesize 1V209-K_2_, 2 equivalents of 4-((6-Amino-2-(2-methoxyethoxy)-8-oxo-7,8-dihydro-9H-purin-9-yl)methyl)benzoic acid (Ambeed, Arlington Heights, IL), also known as 1V209, was preactivated with HATU (1.95 eq.) and DIEA (3 eq.) and allowed to react with the resin overnight. Near-infrared fluorescent peptides were made by coupling 2 equivalents *N*-hydroxysuccinimide (NHS) ester-functionalized ATTO 647N (ATTO-TEC, Siegen, Germany) to the N-terminus with 4 equivalents of DIEA overnight protected from light. TAMRA-E_2_ was prepared identically to TAMRA-K_2_ but with substituting the lysine residues for glutamic acid. In the case of the standard K_2_ peptide, the N-terminus was acetylated with the addition of acetic anhydride (200 eq.) and DIEA (75 eq.) in dichloromethane (DCM) two times for 45 min. In all cases, peptides were cleaved from the resin by adding a cleavage cocktail composed of 90:2.5:2.5:2.5:2.5 trifluoroacetic acid (TFA):anisole:triisopropylsilane:ethylendithiol:water for 3 h at ambient temperature. TFA was evaporated off with a stream of nitrogen gas to a final volume of 1 mL. The peptide was then precipitated with cold diethyl ether.

Crude peptide was purified by high-performance liquid chromatography (HPLC) on an XBridge Protein BEH C4 OBD Prep column (Waters Corporation, Milford, MA) using a solvent system of Milli-Q (MQ) water with 0.05% TFA (solvent A) and acetonitrile with 0.05% TFA (solvent B). Peptides were dissolved for injection at 10-15 mg/mL in 80% solvent A and 20% solvent B. Peptides were purified using a 3% solvent B/min gradient from 5% to 50% solvent B. The same HPLC method was used to purify TAMRA-E_2_ but with a basic pH 8.5 solvent system composed of MQ water with 4 mM acetic acid and 5 mM ammonium hydroxide (solvent A) and acetonitrile with 4 mM acetic acid and 5 mM ammonium hydroxide (solvent B). The purity and identity of the peptides were confirmed by matrix-assisted laser desorption/ionization mass spectrometry (MALDI MS) and ultra-performance liquid chromatography (UPLC) (Figures S1-6,8-10).

### Ovalbumin Fluorophore Labeling

OVA (Millipore Sigma) was labeled with NHS ester-functionalized DyLight 755 (Thermo Scientific, Waltham, MA) using the protocol provided by the supplier. In brief, the protein was dissolved in 0.2 M sodium bicarbonate buffer (pH 8.5) at a concentration of 10 mg/mL. NHS ester-functionalized DyLight 755 was slowly added in stoichiometric amounts to the protein solution. The reaction proceeded at ambient temperature in the dark for 1.5 h. The material was then purified by size exclusion chromatography (SEC) using a HiLoad Superdex 200 pg preparative SEC column (Cytiva, Marlborough, MA) to remove unreacted dye and endotoxins.

### Endotoxin Testing

All materials were verified to be free of endotoxins using the HEK-Blue mTLR4 reporter cell line (InvivoGen, San Diego, CA) using the protocol provided by the supplier. In brief, 20 μL of sterile 1 mg/mL peptide or OVA in endotoxin-free water (Cytiva) was placed in the bottom of a 96-well tissue culture-treated microplate. An endotoxin bacterial LPS standard curve was plated over a range of 10-0.15 EU/mL. HEK-Blue mTLR4 cells (180 μL) at a density of 2.2 x 10^5^ cells/mL were then placed on top of the samples and allowed to incubate at 37 °C and 5% CO_2_ for 20-24 h. Supernatant (20 μL) was then removed from each well and added to 180 μL of QUANTI-Blue detection buffer (InvivoGen). The detection solution was incubated at 37 °C for 0.25-6 h until the lowest standard had visibly turned blue. The absorbance at 630 nm was then read using a microplate reader (Tecan, Männedorf, Switzerland), and the endotoxin content of each sample was determined using the standard curve. All materials were confirmed to contain less than 0.25 EU per mg of material (i.e., OVA or peptide).

### TLR7 Activity Assay

The ability of 1V209-modified peptides to activate TLR7 was assessed using HEK-Blue mTLR7 reporter cells (InvivoGen) using the standard protocol provided by the supplier. Sterile stock solutions of 1V209 or 1V209-conjugated peptides were prepared in endotoxin-free water and diluted to 100 μM. The concentration of each sample was confirmed using a Cary 60 UV-Vis (Agilent Technologies, Santa Clara, CA) by measuring the absorbance at 280 nm. These stock solutions were then added (20 μL) to a TC-treated 96-well plate and HEK-Blue mTLR7 cells (180 μL) were then plated on top of the samples at a concentration of 2.2 x 10^5^ cells/mL and incubated at 37 °C and 5% CO_2_ for 20-24 h. When interrogating the role of cathepsin B in the processing of the TLR7 agonist peptides, the wells also contained 30 μM of the broad-spectrum cysteine protease inhibitor E64D (MedChem Express, Monmouth Junction, NJ). After incubation, 20 μL of supernatant from each well was added to 180 μL of QUANTI-Blue detection media and incubated at 37 °C for 6 h. TLR7 activity was determined by reading the absorbance of the detection solutions at 630 nm using a microplate reader. The absorbance of each sample was normalized to the mean of the 1V209 samples for each experiment.

### Cell Uptake Assay

Tetramethylrhodamine (TAMRA), a small-molecule fluorescent dye, was conjugated to the N-terminus of K_2_ and the negatively charged MDP E_2_ (TAMRA-EE(SL)_6_EE-Am) as described above. Both peptides, along with a TAMRA-only control, were dissolved in sterile, double-distilled water at an approximate concentration of 1 mg/mL and the pH of the solution was adjusted to be between 7 and 8. The absorbance at 540 nm of each solution was used to subsequently standardize the concentration to 100 μM, assuming a molar extinction coefficient of 90,000 M^-1^cm^-1^ for TAMRA. These stocks were filter-sterilized and stored at 4 °C prior to use.

HEK293 cells were seeded at a density of 1.5 x 10^5^ cells/well in a 48-well microplate containing complete media consisting of Dulbecco’s Modified Eagle Medium (DMEM) with 10% fetal bovine serum (FBS) and 1% penicillin-streptomycin, and allowed to recover overnight. The next day, the cells were washed once with phosphate-buffered saline (PBS) and 180 μL of Opti-MEM was added to each well. To this, 20 μL of 100 μM peptide or TAMRA dye stock solution (final concentration = 10 μM) or 20 μL of Opti-MEM was added and the plate was incubated at 37 °C for 4 h. Next, the cells were washed twice with PBS, dissociated using 20 μL of 0.25% Trypsin, 0.1% EDTA solution, and resuspended in 180 μL Dulbecco’s PBS containing Mg^2+^ and Ca^2+^ supplemented with 0.5% FBS and 0.1% 4’,6-diamidino-2-phenylindole (DAPI). Samples were passed through a 70 μm cell strainer and kept on ice prior to performing flow cytometry using a SA3800 instrument (Sony Biotechnology, San Jose, CA). Subsequent analysis was conducted using FlowJo software (FlowJo LLC, Ashland, Oregon).

### Vaccine and Hydrogel Preparation

MDP hydrogels were prepared by making 40 mg/mL peptide stock solutions in ultrapure endotoxin-free water (Cytiva). Once the peptide was completely dissolved, the solution was diluted to 20 mg/mL peptide with 2X Hank’s balanced salt solution (HBSS) containing 400 μg/mL endotoxin-free OVA. The gels composed of a 50:50 mixture of K_2_ and 1V209-K_2_ were made by first preparing two peptide stock solutions each at 40 mg/mL and then mixing the two solutions to make one solution with 20 mg/mL K_2_ and 20 mg/mL 1V209-K_2_. This combined stock solution was then diluted 1:1 with the 2X OVA stock solution in buffer to form the hydrogel. To prepare the gel with 100 μM 1V209-K2, 1V209-K_2_ was dissolved at 200 μM in water and then this solution was used to prepare a stock solution of 40 mg/mL K_2_. Gels from this solution were prepared identically as described above. To make hydrogels without OVA, the same protocol was followed, but OVA was not included. Solutions were then incubated at 4 °C in the dark overnight to allow for complete gelation. Alum samples were prepared by performing a 1:1 dilution of 2% Alhydrogel® adjuvant (InvivoGen) with stock solutions of 400 μg/mL endotoxin-free OVA. OVA-only vaccines were prepared by performing a 1:1 dilution of the OVA stock solution in 2X HBSS with the same volume of endotoxin-free water. 1V209 was prepared in 150 mM NaCl at a concentration of 100 μM with 200 μg/mL OVA, which was near the limit of solubility for 1V209. All vaccine formulations had a final OVA concentration of 200 µg/mL.

### Rheology

Oscillatory rheology measurements were performed using an AR-G2 rheometer (TA Instruments, New Castle, DE). Hydrogel samples (75 μL) without OVA were transferred to the instrument with a truncated 200 μL pipet to minimize shear and were equilibrated at 1 rad/s and 1% strain at ambient temperature for 30 min. After equilibration, frequency sweep data were collected over the range of 0.1-10 rad/s at a constant 1% strain. After a two-minute equilibration period, amplitude sweeps were then performed at an oscillatory frequency of 1 rad/s over a strain range of 0.1-200%. Lastly, shear recovery experiments were conducted by subjecting the hydrogels to 200% strain for 1 min immediately after the amplitude sweep. Recovery of the storage modulus (G’) and loss modulus (G’’) were then monitored for 10 min at the starting conditions. The percent recovery of the hydrogel was determined by dividing the final storage modulus values at the end of the 10 min by the value obtained at the end of the initial two-minute equilibration.

### Fourier Transform Infrared Spectroscopy (FTIR)

Attenuated total reflectance FTIR spectra were collected using a Nicolet^TM^ iS20 FTIR spectrometer (Thermo Fisher Scientific). The atmosphere in the instrument was replaced with a steady stream of nitrogen to avoid capturing atmospheric molecular vibrations. Hydrogel samples (10 μL) without OVA were plated on the instrument and allowed to dry completely. Absorbance spectra were obtained as an accumulation of 30 measurements and background subtracted. Spectra were area-normalized from 1575-1705 cm^-1^ using Python 3.0 to focus on the Amide I band.

### *In Vivo* Fluorescence Imaging

All animal studies were conducted in accordance with an IACUC-approved protocol (22-246) in the Animal Resource Facility at Rice University. One day before beginning experiments, MDP hydrogels were prepared with 200 μg/mL DyLight 755-labeled OVA and 300 μg/mL Atto647N-labeled K_2_ peptide. All other vaccine injections were prepared as described above. Samples (50 μL) were loaded into 31-gauge insulin syringes and stored overnight protected from light at 4 °C. Immediately before injection, the background tissue autofluorescence in the expected wavelengths (Ex/Em 640/700 nm and 750/800 nm) of each mouse was acquired with an In Vivo Imaging System (IVIS) small animal imager (PerkinElmer, Waltham, MA). The background signal was subtracted from all subsequent measurements. SKH1-Elite mice (Charles River Laboratories, Wilmington, MA) then received subcutaneous injections of various vaccine formulations in the hind flank and the fluorescence intensity of OVA (Ex/Em 640/700 nm) and K_2_ peptide (Ex/Em 750/800 nm) was monitored until it became undiscernible from the background. Release was calculated by drawing equally sized rectangular regions of interest around the injection site and dividing the observed total radiant efficiency by the maximum total radiant efficiency observed on the first day of the experiment. OVA release and peptide clearance were measured by monitoring the decrease in fluorescence at the site of injection at their respective wavelengths over time.

### Serum Collection and Antibody Titers

One day before administering the vaccine, naïve serum was collected by bleeding SKH1-Elite mice from the submandibular vein and collecting 200 μL of the whole blood in clotting microvettes (Sarstedt, Nümbrecht, Germany). Serum was then extracted from the whole blood by centrifuging the microvettes at 10,000 rcf for 10 min at 4 °C and stored at −20 °C until further use. Serum was collected every two weeks after vaccination and stored in the same manner through week 12.

The anti-OVA IgG titers in serum were measured by ELISA. Nunc Maxisorp 96-well plates (ThermoFisher Scientific) were coated with 100 μL of 1 μg/mL OVA in a 100 mM pH 9.6 carbonate– bicarbonate buffer overnight at 4 °C. Coated plates were then washed three times with PBST (0.5% Tween 20 in 1X phosphate buffer) using a 96-well plate automatic BioTek Microplate Washer 405 (Agilent) and subsequently blocked with Blotto non-fat dry milk (Rockland Immunochemicals, Limerick, PA) dissolved at 50 mg/mL in PBST (blocking solution) for 2 h at ambient temperature on an orbital shaker. The blocking solution was then removed, and serum samples were then loaded into the plate with an initial ten-fold dilution in blocking solution followed by serial two-fold dilutions. Next, the plates were incubated at ambient temperature for 2 h on an orbital shaker and washed 3 times with PBST. Horseradish peroxidase (HRP)-conjugated rabbit anti-mouse IgG (Jackson ImmunoResearch, West Grove, PA) was diluted to 800 ng/mL in blocking solution and then 100 μL was added to each well and incubated for another 2 h at ambient temperature. The plates were then washed five times with PBST before being developed with 100 μL SureBlue TMB substrate solution (SeraCare Life Sciences Inc., Milford, MA) for 3 min. Development was stopped with the addition of 100 μL of 1M sulfuric acid and the signal in each well was measured as the difference in absorbance at 450 nm and the reference wavelength 640 nm on a microplate reader. Antibody titers were then calculated by determining the lowest serum dilution where the signal was at least two-fold higher than the background signal from naïve serum.

Antibody subclass titers were determined using the same method as above, substituting HRP-conjugated goat anti-mouse IgG1, IgG2a, IgG2b, and IgG3 antibodies (Jackson ImmunoResearch) as the secondary antibody. The initial sample dilution was 64-fold due to serum volume limitations, followed by serial two-fold dilutions. All subclasses were developed for 3 min except for IgG3, which was developed for 10 min.

### Flow Cytometry

OVA was prepared with MDP hydrogels, alum, 1V209, or HBSS and administered in SKH1-Elite mice (n=4 per group) as described above. Seven days after injections, mice were euthanized using carbon dioxide asphyxiation and their spleens were collected. Each spleen was placed into a 100 µm cell strainer (Fisher Scientific, Hampton, NH) in a 50 mL centrifuge tube pre-wetted with sterile HBSS and homogenized aggressively with the back of a sterile 10 mL syringe plunger. The plunger and filter were rinsed into the centrifuge tube with an additional 3 mL of HBSS. This solution was then pipetted onto a 70 µm cell strainer (Fisher Scientific) in a new 50 mL centrifuge tube to remove particulates. The flow-through of this filter was pipetted through a 35 µm cell strainer into a flow tube (Corning, Corning, NY). Tubes were then centrifuged at 250 rcf at 4 °C for 5 min and the supernatant was removed. Cell pellets were resuspended in 1 mL of cold ACK lysis buffer (Quality Biological, Gaithersburg, MD) for 30 sec, then diluted into 10 mL of cold sterile PBS. Samples were centrifuged at 250 rcf at 4 °C for 5 min and the supernatant was removed. Cell pellets were then resuspended in 1 mL of flow buffer and cell counts were determined using a Countess II FL (Invitrogen). Samples were diluted to 2 x 10^7^ cells per mL and 50 µL of each sample was added to 5 mL tubes (Eppendorf, Hamburg, DE), to constitute 1 million cells per flow sample.

After preparing cell suspensions, 28 µL of flow buffer, 2 µL of Fc block, and 10 µL of Brilliant Stain Buffer (Becton Dickinson, Franklin Lakes, NJ) were added to each tube. Anti-H-2K^b^-SIINFEKL antibody (BioLegend, San Diego, CA) was added to all samples, then incubated on ice in the dark for 45 min. The remaining antibodies were then added (Table S1) and samples were incubated on ice in the dark for 30 min. After adding 1 mL of cold PBS, the samples were centrifuged at 200 rcf for 7 min at 4 °C and the supernatant was removed. Cells were resuspended in 1 mL of PBS containing 1 µL of LIVE/DEAD Fixable Yellow Stain (Thermo Fisher Scientific). Cells were incubated on ice in the dark for 30 min, then centrifuged at 200 rcf for 7 min at 4 °C. The supernatant was removed and cells were resuspended in 1 mL of flow buffer, passed through a 35 µm cell strainer into a flow tube, and kept on ice for < 2 h before analysis on a Sony SA3800 Flow Cytometer. Results were processed using FlowJo software.

When testing for MHC I H-2K^b^, five SKH1-Elite mice from different litters and of different ages, ranging from one to six months, were selected to ensure maximum variability of potential MHC I haplotype. One C57BL/6 mouse was used as a positive control and one BALB/c mouse was used as a negative control. The anti-mouse H-2K^b^ antibody used was confirmed to be negative for cross-reactivity with other haplotypes. The protocol used was identical to the one described above up until the addition of the first antibody; the only stain used was anti-mouse H-2K^b^ conjugated to APC (BioLegend). The samples were incubated on ice in the dark for 45 min before 1 mL of cold PBS was added and the samples were centrifuged at 200 rcf for 7 min at 4 °C. The supernatant was removed and cells were resuspended in 1 mL of flow buffer, passed through a 35 µm cell strainer into a flow tube, and kept on ice for 20 min before analysis on a Sony SA3800 Flow Cytometer. Results were processed using FlowJo software.

### Histology

One week after vaccination, the skin surrounding the injection site of SKH1-Elite mice was harvested, pinned to wax, and fixed in 10% formalin overnight at 4 °C. The formalin was then drained and replaced with a solution of 30% sucrose in DI water, in which the samples were stored overnight at 4 °C. Skin samples were then patted dry and frozen in Optimal Cutting Temperature (OCT) compound on dry ice and stored at −80 °C until being sectioned at 5 µm on a cryostat.

Sections were stained with Scytek Laboratories (Logan, UT) Hematoxylin and Eosin (H&E) Stain Kit according to the manufacturer’s protocol with slight modifications. Slides were stained with hematoxylin for 15 sec, washed in two changes of deionized water, then treated with Bluing Reagent for 15 sec and again washed in two changes of deionized water. Slides were then dipped in eosin for 2 sec and rinsed in ethanol. Slides were dehydrated in 3 changes of ethanol and cleared in 3 changes of xylene before mounting using Shandon Mount (Epredia, Kalamazoo, MI) and imaging on an EVOS M5000 Microscope (Thermo Fisher Scientific).

Additional sections were stained with Masson Trichrome Stain Kit (Epredia) according to the manufacturer’s protocol, adapted for use on frozen sections. Briefly, slides were incubated in Bouin’s Fluid at 56 °C for 1 h and rinsed in tap water, then placed in Weigert’s Iron Hematoxylin for 3.5 min. Slides were rinsed, stained in Biebrich Scarlet-Acid Fuchsin for 2.5 min, then rinsed again. Finally, slides were stained with Aniline Blue for 30 sec, dipped in 1% acetic acid, and rinsed. Slides were dehydrated in 3 changes of ethanol and cleared in 3 changes of xylene before mounting using Shandon Mount and imaging on an EVOS M5000 Microscope.

### Immunohistochemistry

Tissue sections collected for histology were also stained for F4/80, CD11c, and TNF-α by immunohistochemistry. Slides were incubated at ambient temperature in blocking solution (SuperBlock Buffer, Thermo Scientific) for 30 min. Biotinylated anti-F4/80, anti-CD11c, and anti-TNF-α antibodies (Table S2) were diluted in blocking solution and incubated on slides overnight at 4 °C. Slides were treated with 3% hydrogen peroxide for 10 min at ambient temperature to remove endogenous peroxidases, then washed in PBS and incubated with streptavidin-HRP (BioLegend) for 1 h at 4 °C. Slides were washed and DAB substrate (Becton Dickinson) was added for 15 min at ambient temperature. Samples were then washed in PBS and stained with hematoxylin (from H&E protocol above). Finally, slides were cleared in three changes of xylene, mounted in Shandon Mount, and imaged on an EVOS M5000 Microscope.

### Statistics

Single-group comparisons were performed by unpaired Student’s t-tests. Multiple group comparisons were calculated by ordinary one- or two-way ANOVA with Tukey’s multiple comparisons test. All statistical analyses were performed in GraphPad Prism 10. Statistical significance in all figures is denoted with asterisks as follows: *p < 0.05; **p < 0.01; ***p < 0.001; ****p < 0.0001. All data is reported as the mean ± standard deviation and error bars indicate one standard deviation from the mean.

## Results and Discussion

### *In Vitro* Activity of TLR7-agonist-modified MDPs

Since previous studies have shown that linker length and the presence of enzymatic cleavage sites covalently attaching TLR7 agonists to biomaterials can impact the adjuvancy of the composite,^[17,38]^ we developed a small library of five 1V209 and K_2_ MDP conjugates (Figure 2A) with varying linker length by including a PEG spacer and a cathepsin B-cleavable peptide sequence (GFLG) to screen for the optimal peptide design for TLR7 agonism (Figure S1-6). Both the cathepsin B enzyme and TLR7 are expressed in the endosomes of APCs, and the GFLG sequence has been extensively used to liberate payloads in the presence of this enzyme.^[39]^ The TLR7 agonist activity of these peptides was assessed using HEK-Blue TLR7 reporter cells and compared to equimolar amounts of unconjugated 1V209. We observed that conjugating 1V209 to MDPs affords an over 200-fold improvement in the solubility of the small-molecule agonist in water. The maximum solubility of the 1V209 small molecule in water is approximately 100 μM whereas all of the 1V209-conjugated peptides studied can be dissolved at > 20 mM. Thus, all materials were tested at a concentration of 10 μM so the unconjugated 1V209 small molecule could be easily dissolved.

**Figure 2.**
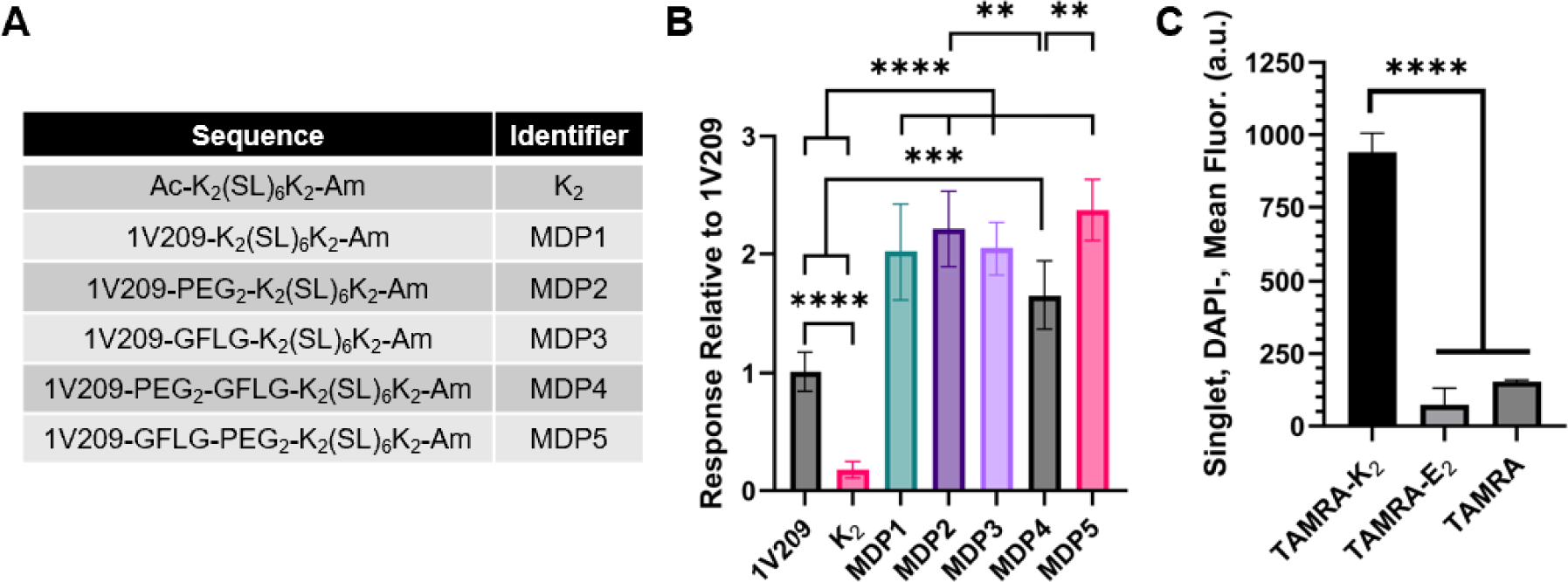
A) The amino acid sequences of each synthesized MDP. B) Measurements of TLR7 agonist activity in (A), normalized to the activity of unconjugated 1V209. C) Measured cell uptake of TAMRA-conjugated MDPs, showing enhanced uptake when using positively charged K_2_ compared to negatively charged E_2_ or TAMRA alone.

All five 1V209-modified peptides significantly enhanced TLR7 agonism compared to the small molecule alone (Figure 2B). The differences in TLR7 agonist activity between MDP1, MDP2, MDP3, and MDP5 were not statistically significant while MDP2 and MDP5 were both significantly different from MDP4. As expected, K_2_, which lacks 1V209, did not display meaningful TLR7-mediated responses. To assess if the cathepsin linker was responsible for any of the observed differences between the 1V209-modified MDPs, reporter cells were again incubated with the five MDPs and 1V209 in the presence of 30 μM of the broad-spectrum cysteine protease inhibitor E64D, which blocks the activity of several proteases, including cathepsin B. The presence of the inhibitor significantly reduced the TLR7 agonist activity of all the peptides except for MDP4, suggesting that protease activity in the endosome contributes to TLR7 agonism by directly breaking down K_2_ to liberate 1V209 and not through cleavage of the GLFG motif (Figure S7). Together, these results suggest that linkers and cleavage sites are not required to retain the activity of 1V209 and that attaching it directly to the N-terminus of K_2_ does not undermine its activity. Therefore, MDP1, which we will henceforth refer to as 1V209-K_2_, was used for all subsequent studies.

Interestingly, the initial peptide screen showed that linking 1V209 to K_2_ increased the agonist activity of the small molecule approximately two-fold. We hypothesized that K_2_ may improve the cell uptake of 1V209 because it is amphiphilic and positively charged, resembling known cell-penetrating peptides.^[40,41]^ To test this hypothesis, a cell uptake assay was performed using K_2_ modified at the N-terminus with the small-molecule fluorescent dye tetramethylrhodamine (TAMRA). A modified version of the negatively charged MDP E_2_ (TAMRA-E_2_) where all the lysine residues of K_2_ are replaced with glutamic acid residues (Figure S8-9) was used as a comparator. TAMRA-K_2_ improved cell uptake by a factor of six compared to TAMRA and by a factor of ten compared to TAMRA-E_2_ (Figure 2C). These data suggest that the improved TLR7 agonist activity of 1V209-K_2_ may be attributed to the positive charge of the K_2_ MDP that improves cell penetration.

### Materials Characterization of 1V209-K_2_ Supramolecular Hydrogels

Unmodified MDPs are known to form antiparallel β-sheets that self-assemble into a network of nanofibers.^[33]^ To ensure that the covalent addition of 1V209 did not significantly interfere with self-assembly, FTIR was employed to assess the secondary structure of 1V209-K_2_. The FTIR spectra of K_2_, 1V209-K_2_, and a 50:50 blend of K_2_:1V209-K_2_ all showed two peaks at approximately 1620 cm^-1^ with an additional peak at 1695 cm^-1^, indicating the formation of antiparallel β-sheets^[42]^ (Figure 3A). The only differences observed between the groups were minor, occurring at approximately 1675 cm^-1^, which can be attributed to the carbonyl stretch of residual TFA from HPLC purification.^[43]^

**Figure 3.**
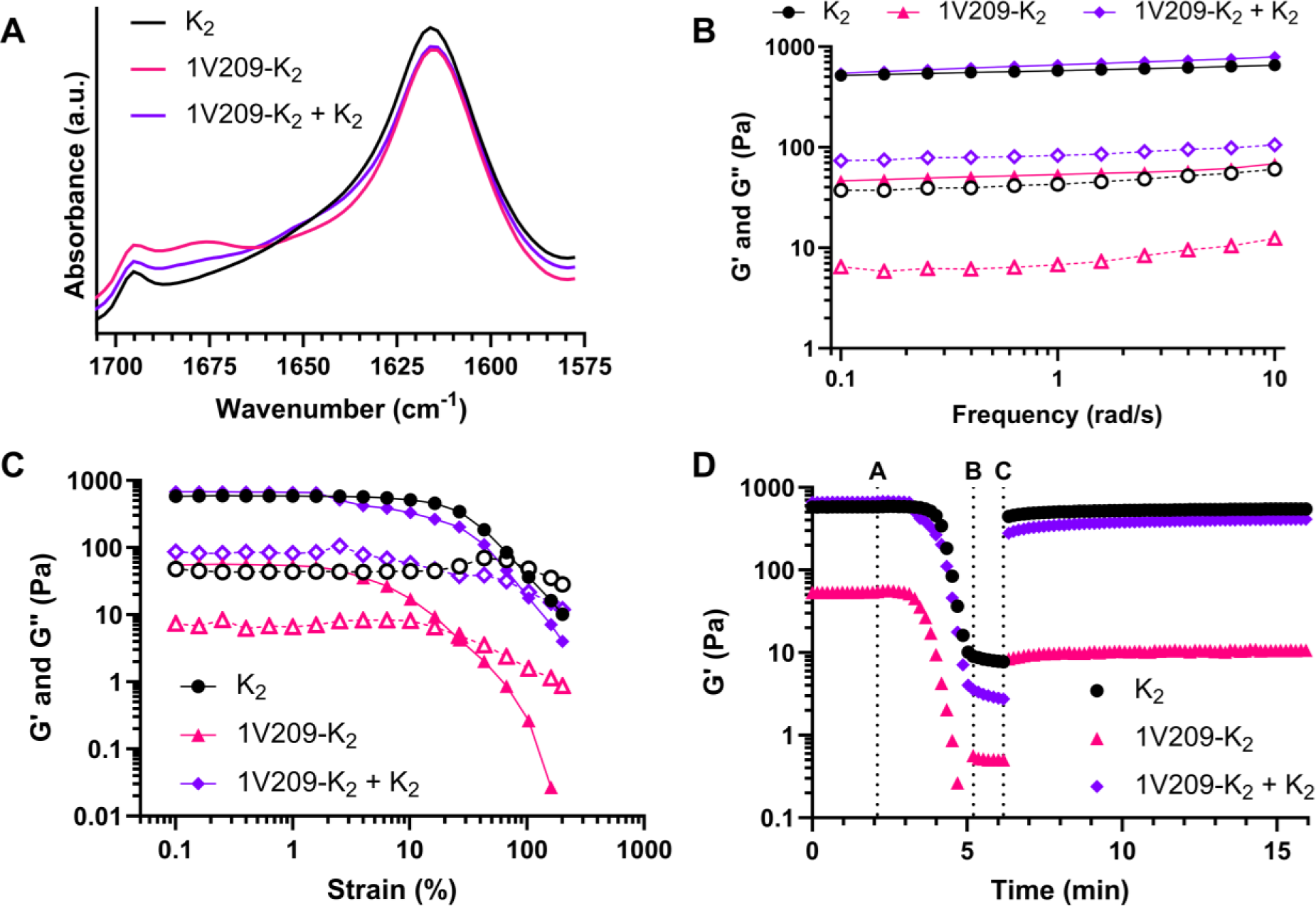
A) FTIR spectra of K_2_, 1V209-K_2_, and a 50:50 (w/w) mixture of K_2_ and 1V209-K_2_ (K_2_ + 1V209-K_2)_ indicating that all samples form antiparallel β-sheets as evidenced by the presence of peaks at ∼1620 cm^-1^ and 1695 cm^-1^. B) Frequency sweeps of the three hydrogels from 0.1 – 10 rad/s showing that the behavior of all three gels is frequency-independent over this range. Filled symbols and solid lines indicate the storage modulus (G’) and empty symbols and dotted lines indicate the loss modulus (G”). C) Amplitude sweeps from 0.1 – 200% strain demonstrate that all three gels exhibit shear thinning and that pure 1V209-K_2_ has the smallest linear viscoelastic regime. Filled symbols and solid lines indicate G’ and empty symbols and dotted lines indicate G”. D) Shear recovery experiments showed that 1V209-K_2_ and K_2_ + 1V209-K_2_ are self-healing after the amplitude sweep experiment (between lines A and B) and after being subjected to 1 min of deformation of 200% strain (between lines B and C).

After the secondary structure of each peptide self-assembly was confirmed, the ability of 20 mg/mL 1V209-K_2_ to form a hydrogel in 1X HBSS was assessed by oscillatory rheology and compared to 20 mg/mL K_2_ and a 50:50 (w/w) blend of K_2_:1V209-K_2_ (K_2_ + 1V209-K_2_) dissolved at a final total peptide concentration of 20 mg/mL. Upon the addition of HBSS, the pure 1V209-K_2_ hydrogel changed from transparent to cloudy and formed a weak gel by visual inspection. In contrast, pure K_2_ formed a completely transparent gel and K_2_ + 1V209-K_2_ was mostly clear with some minor turbidity when HBSS was added to trigger gelation. These qualitative assessments of gel formation agreed with subsequent frequency sweep rheological data, which showed that K_2_ and 1V209-K_2_ + K_2_ had very similar G’ in the range of 500-600 Pa while 1V209-K_2_ alone had a G’ about an order magnitude lower around 40 Pa, which is similar to the G” of the two other gels. Pure K_2_ formed the strongest gel, as indicated by having the highest G’ and the lowest G”. The moduli of all three gels were largely frequency-independent between 0.1 and 10 rad/s (Figure 3B). Amplitude sweeps over the range of 0.1 to 200% strain showed that K_2_ alone is linearly viscoelastic up to 20% strain, after which its G’ drops rapidly. In comparison, 1V209-K_2_ is only linearly viscoelastic up to 2% strain. 1V209-K_2_ + K_2_ appeared to behave like a combination of the two homogenous hydrogels with two distinct domains where it is largely able to resist deformation. Up to 2% strain, the gel was linearly viscoelastic, but between 2% and 20% the drop in G’ was very slow (Figure 3C). Despite the differences in the linear viscoelastic regimes of 1V209-K_2_ + K_2_ and K_2_ alone, the hydrogel character was lost (G” > G’) in both materials at about 100% strain while the pure 1V209-K_2_ hydrogels exhibited liquid-like behavior starting at 30% strain.

Amplitude sweeps of all three hydrogels indicated that they are all shear-thinning viscoelastic hydrogels capable of flowing through small-bore needles upon injection. To confirm that these materials are self-healing after deformation, a shear recovery experiment was conducted after the amplitude sweeps by holding the gels at 200% strain for 1 min and then reducing the strain back to 1% for 10 min to watch the recovery in G’ (Figure 3D). The K_2_ hydrogel completely recovered its pre-deformation G’ almost immediately, while the 1V209-K_2_ + K_2_ blended hydrogel recovered 41% of its initial G’ 10 sec after the high shear force was abated and 61% by the end of the 10 min equilibration period. In contrast, pure 1V209-K_2_ hydrogels only recovered 19% of their initial G’. Taken together, rheological testing indicated that blending 1V209-K_2_ with K_2_ yields a relatively strong shear-thinning and self-healing hydrogel that has properties more suitable for biomedical application than hydrogels composed of 1V209-K_2_ alone.

### *In Vivo* Antigen Release and Hydrogel Persistence

Without controlled release, small-molecule TLR7 agonists rapidly diffuse from the site of injection and are quickly cleared from the body—typically within a few hours.^[44]^ By covalently conjugating 1V209 to K_2_, we hypothesized that TLR7 agonist activity would be prolonged as the modified peptides slowly disassemble and are cleared. To determine hydrogel degradation kinetics and investigate if the inclusion of 1V209 alters hydrogel clearance, K_2_ and 1V209-K_2_ + K_2_ were each doped with 1.5% by mass of K_2_ conjugated to the fluorophore Atto647N at the N-terminus (Figure S10). These materials were injected subcutaneously into the flanks of SKH1-Elite mice and fluorescence at the site of injection was monitored over time to determine the clearance rate. Hydrogels composed of K_2_ and 1V209-K_2_ + K_2_ exhibited statistically similar degradation profiles *in vivo*, each losing approximately half of their initial fluorescence within 20-25 days and persisting for over 85 days (Figure 4A). Although this experiment does not directly measure the persistence of 1V209-K_2_, mice that received 1V209-K_2_ + K_2_ injections exhibited visible signs of mild inflammation at the injection site throughout the 85-day study while no inflammation was observed when administering K_2_ alone, suggesting that 1V209-K_2_ remained at the site of injection for the duration over which fluorescence was detected.

**Figure 4.**
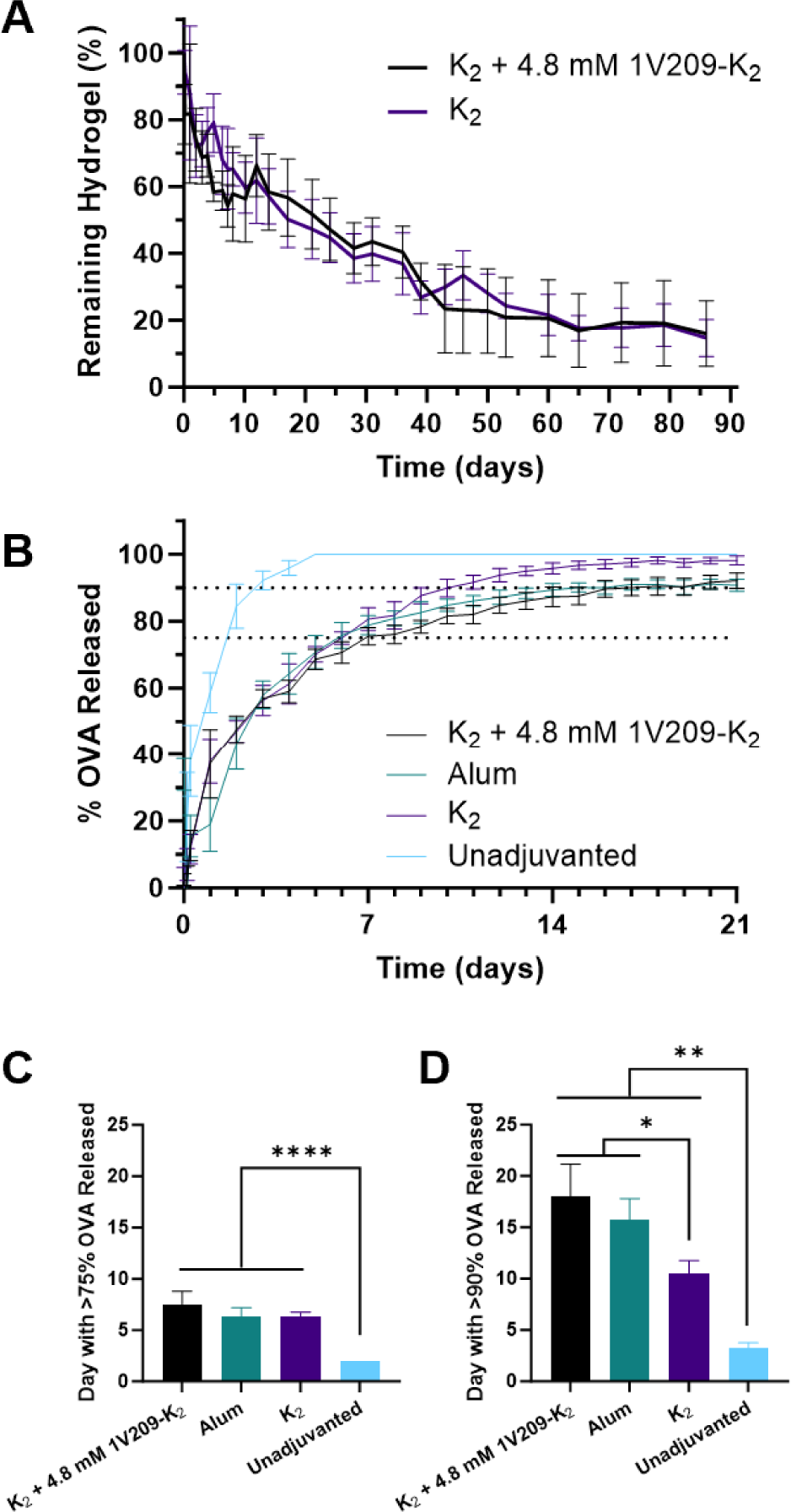
A) *In vivo* degradation of MDP hydrogels as measured by clearance of Atto647N-K_2_, showing that the inclusion of 1V209-K_2_ has no effect. B) *In vivo* OVA release from the injection site as measured by clearance of DyLight 755-conjugated OVA. Unadjuvanted free OVA clears fully in 5 days, while the three hydrogel groups delay release significantly. C) The timepoint at which the mice in (B) reached greater than 75% OVA release, showing that all three hydrogel groups are not statistically different and significantly greater than the unadjuvanted group. D) The timepoint at which the mice in (B) reached greater than 90% OVA release, showing that K_2_ + 4.8 mM 1V209-K_2_ and alum retained OVA significantly longer than K_2_.

One way that adjuvants like alum improve immune responses to vaccines is by controlling the release of antigen over time.^[45]^ Since these hydrogels persist for long periods, we hypothesized that 1V209-functionalized hydrogels might be able to prolong the release of antigens *in vivo*. To investigate the ability of these gels to act as an antigen depot, we injected hydrogels loaded with fluorescently labeled OVA injected into mice and then measured the decrease in fluorescence at the injection site over time using an an IVIS. OVA without any hydrogel was rapidly cleared from the injection while all hydrogel groups (including MDPs and alum) appeared to retain some OVA through the end of the 21-day study (Figure 4B). All hydrogel groups released OVA significantly slower than the OVA-only injection, releasing 75% of their OVA after 6-8 days instead of within just 2 days (Figure 4C). However, while K_2_ released 90% of encapsulated OVA by day 10, alum and 1V209-K_2_ + K_2_ released the remaining OVA more slowly, releasing 90% over 16 and 18 days, respectively (Figure 4D).

Since the maximum solubility of 1V209 is 0.1 mM, an additional 1V209-K_2_ hydrogel with 0.1 mM 1V209-K_2_ mixed with K_2_ for a total concentration of 20 mg/mL (referred to hereafter as K_2_ + 0.1 mM 1V209-K_2_) was prepared in addition to the 50:50 (w/w) blend of K_2_:1V209-K_2_ hydrogel (referred to hereafter as K_2_ + 4.8 mM 1V209-K_2_). The co-delivery of unconjugated 1V209 slightly increased the OVA clearance rate compared to unadjuvanted OVA; OVA release from hydrogels composed of K_2_ + 0.1 mM 1V209-K_2_ was cleared at a rate similar to alum and K_2_ + 4.8 mM 1V209-K_2_ (Figure S11). These data show that adding 1V209 to K_2_ improves its ability to retain OVA, which may be due to π-π stacking between aromatic residues on the surface of OVA and 1V209 or attractive interactions between 1V209 and hydrophobic amino acids in OVA.

### Histological Analysis of Subcutaneous Hydrogels

SKH1-Elite mice were vaccinated using the protocol described above and tissue in the rear flanks was excised at 1 and 13 weeks after vaccination to study cellular infiltration and fibrous capsule formation. After one week, the injection sites of the soluble groups did not show any obvious signs of drug administration whereas the hydrogels were all visible by eye. H&E staining showed a visible difference in cell infiltration into 1V209-containing hydrogels as compared to those containing only K_2_ (Figure 5, black dashed lines). Masson’s Trichrome staining showed a small and qualitatively similar fibrous capsule around all four gel types (Figure 5, yellow arrows). The thickness of the dermal layer adjacent to the 1V209-containing gels was visibly thicker (Figure 5, green arrows) than the skin adjacent to K_2_ hydrogels or alum, signifying a greater degree of inflammation.

**Figure 5.**
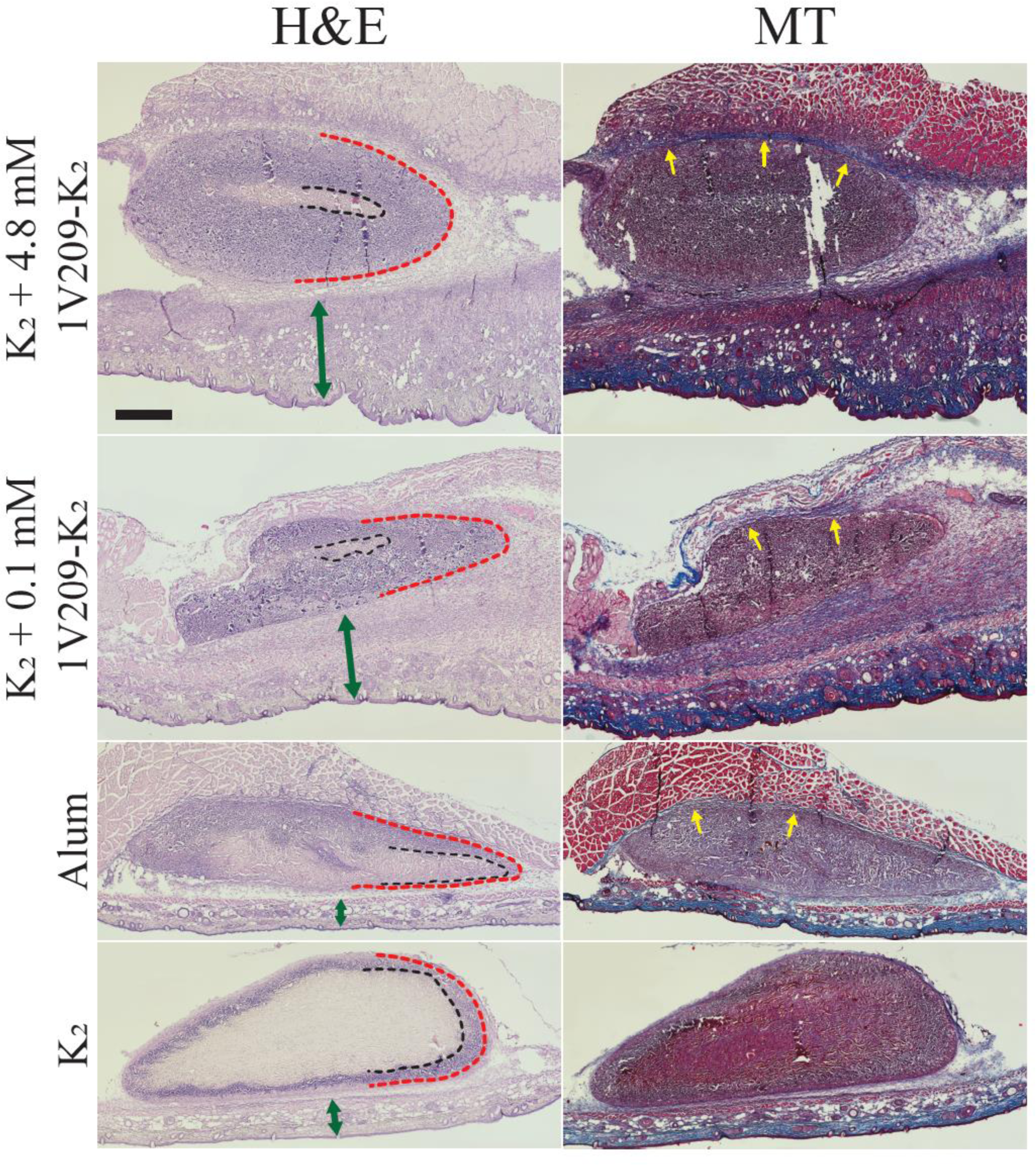
Hematoxylin and eosin (H&E) and Masson’s Trichrome (MT) staining of hydrogels and surrounding tissue excised from mice one week after injection. Inflammation is visible in both 1V209-containing hydrogels, as seen by an increase in skin thickness (green arrows). The dashed red line shows the outside of the hydrogel and the dashed black line shows the edge of cellular infiltration, as identified by the presence of nuclei staining in H&E. Both 1V209-containing hydrogels had visually obvious increases in cellular infiltration compared to K_2_. The 1V209-containing gels and alum all have some fibrous capsule formation (yellow arrows). The scale bar denotes 500 µm.

Thirteen weeks after injection, only a small subset of gels were visible in the subcutaneous space, despite IVIS imaging suggesting that K_2_ and K_2_ + 4.8 mM 1V209-K_2_ gels were still present. We hypothesize that this signal is a result of residual peptide nanofibers persisting at the injection site that are below the gel-forming concentration. Of the tissues containing gels, those containing 1V209-modified peptides developed a qualitatively thicker fibrous capsule at week 13 (Figure 6). Additionally, compared to week 1, these samples showed pervasive cell infiltration and decreased inflammation. In comparison, K_2_ gels had significantly decreased in size.

**Figure 6.**
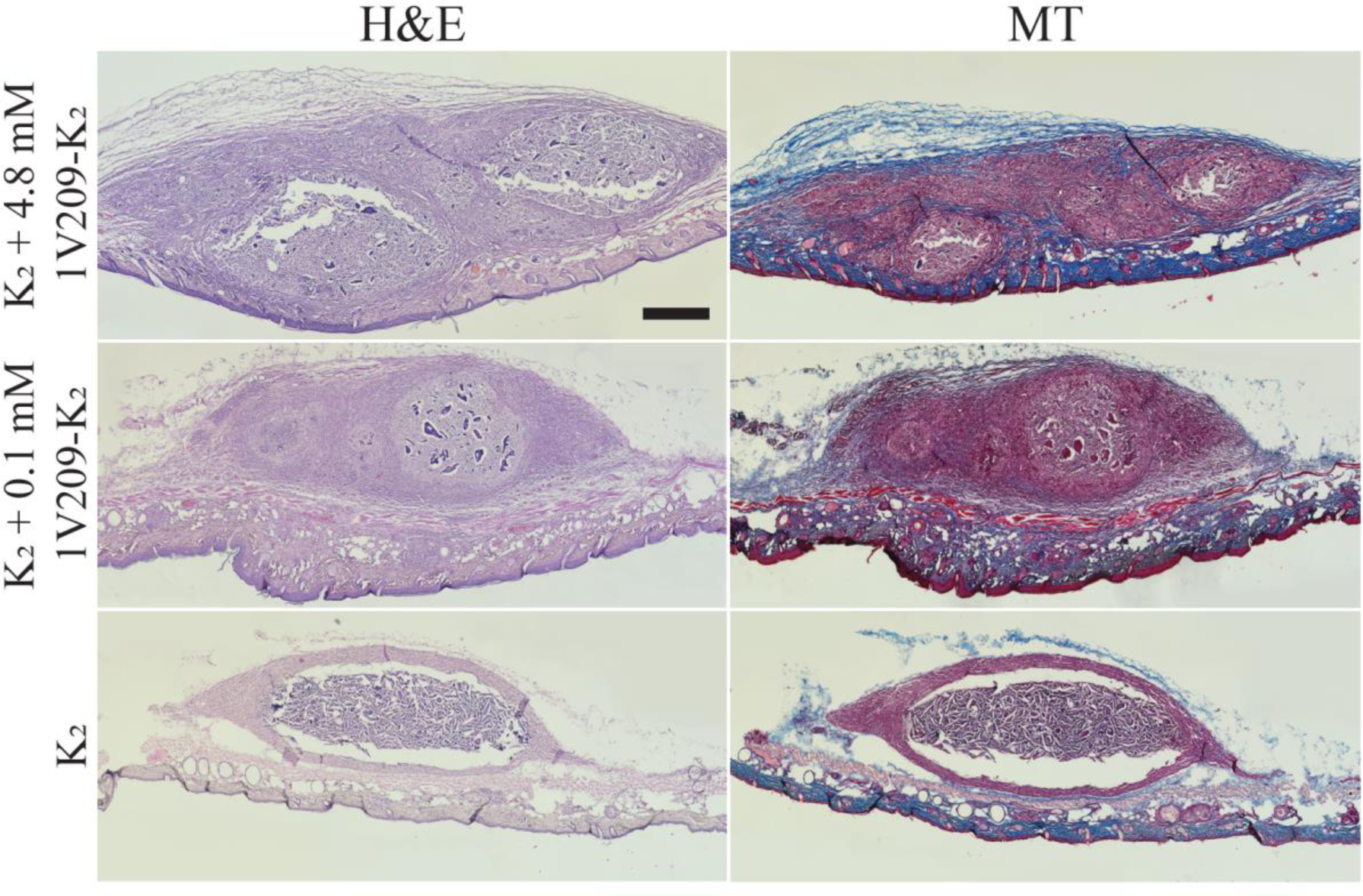
Hematoxylin and eosin (H&E) and Masson’s Trichrome (MT) staining of *in vivo* hydrogels excised 13 weeks after injection. Some inflammation is visible in the K_2_ + 0.1 mM 1V209-K_2_ group, seen as an increase in skin thickness. A blue fibrous capsule is visible in MT staining of both 1V209-containing hydrogels. Alum was not included as the hydrogels were unable to be located in the subcutaneous space. The scale bar denotes 500 µm.

### Immunological Evaluation of 1V209-K_2_ Adjuvancy

The humoral immune response of SKH1-Elite mice to OVA was measured by quantifying OVA-specific IgG antibody titers longitudinally over 12 weeks and IgG subclasses at week 12 in the serum of vaccinated mice. While the antibody titers in mice receiving 0.1 mM 1V209 remained at or near the lower limit of detection throughout the full 12 weeks, the dose-matched K_2_ + 0.1 mM 1V209-K_2_ injections resulted in anti-OVA IgG titers significantly higher than K_2_ after week 4 and not statistically different from alum at any timepoint studied (Figure 7A). OVA delivered in K_2_ + 4.8 mM 1V209-K_2_ hydrogels induced IgG titers that were not statistically different from OVA delivered in K_2_ + 0.1 mM 1V209-K_2_ hydrogels, suggesting that the humoral adjuvancy effect of 1V209-modified K_2_ is not dose-responsive across the two orders of magnitude studied and that local TLR7 activation might already be saturated at the lowest dosage.

**Figure 7.**
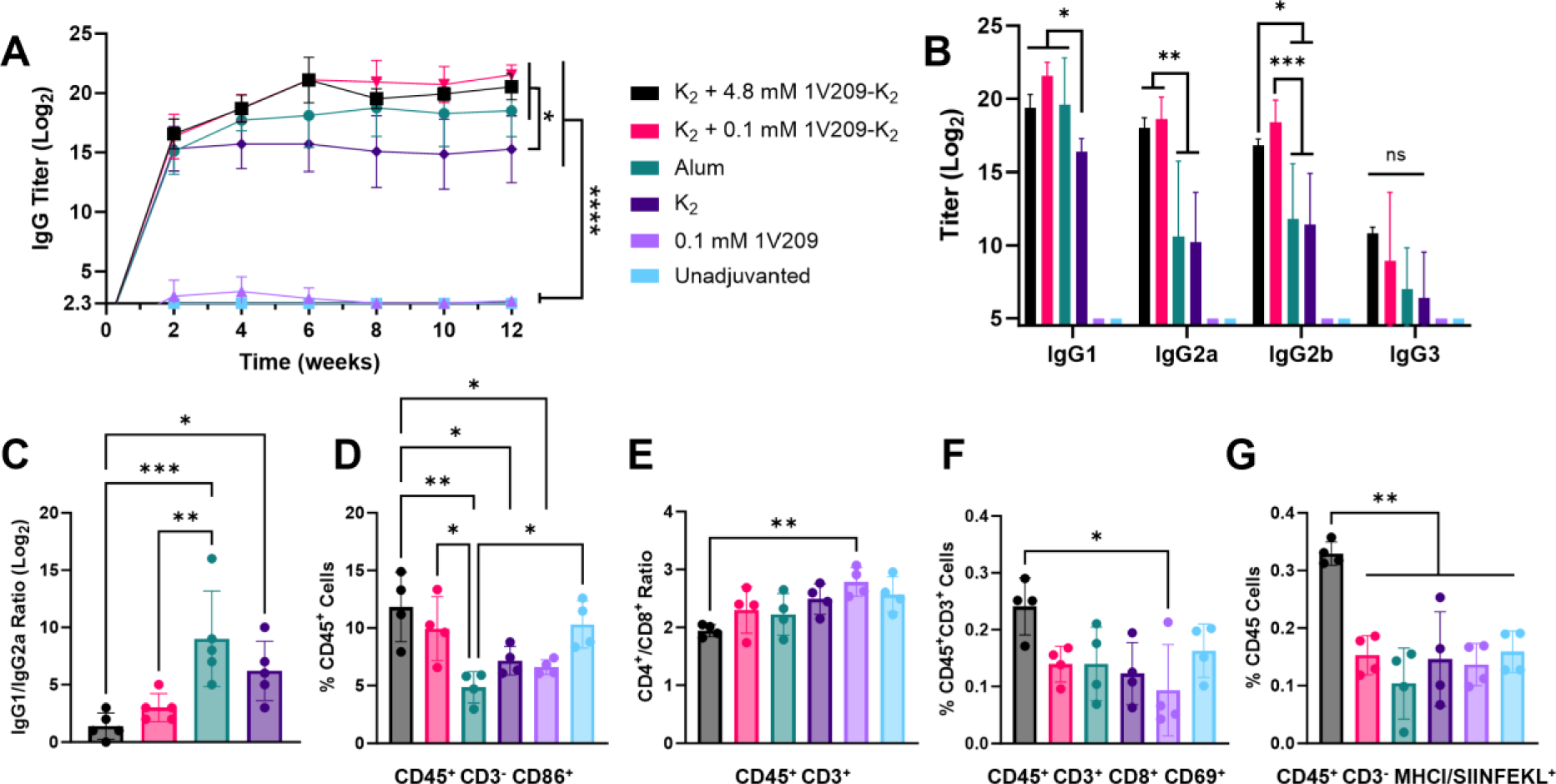
A) Longitudinal anti-OVA IgG titers measured over 12 weeks, with n=5 per group. At the final timepoint, titers for both 1V209-K_2_ groups and alum were significantly greater than those for K_2_, and all four hydrogel groups were significantly greater than the two carrier-free groups, which were both below the limit of detection. The legend in (A) applies to all panels. B) IgG subclasses measured at week 12. All hydrogel groups were significantly greater than the carrier-free groups unless otherwise noted. The 1V209-K_2_ groups were not statistically different for any subclass and both induced significantly higher titers than K_2_ in IgG1, IgG2a, and IgG2b. Both 1V209-K_2_ groups also generated significantly higher titers than alum in IgG2a and IgG2b. K_2_ + 4.8 mM 1V209-K_2_ produced significantly higher IgG3 titers than the two carrier-free groups, but no other groups were statistically different from one another. C) The IgG1/IgG2a ratio at week 12 for all groups with IgG titers above the limit of detection, showing a significant decrease of both 1V209-K_2_ groups below alum and of K_2_ + 4.8 mM 1V209-K_2_ below K_2_. D-G) Flow cytometry data from splenocytes measured one week after vaccination, with n=4 per group. D) Percent of total CD45^+^ lymphocytes that are CD3^-^CD86^+^, signifying non-T lymphocytes expressing a costimulatory activation marker. E) The CD4:CD8 ratio when gated for CD45^+^CD3^+^ T lymphocytes, showing a decrease in the K_2_ + 4.8 mM 1V209-K_2_ group. F) Percent of total CD45^+^CD3^+^ cells co-expressing CD8 and the activation marker CD69, showing an increase in the K_2_ + 4.8 mM 1V209-K_2_ group. G) The percent of total CD45^+^ lymphocytes that are CD3^-^ (signifying likely antigen-presenting cells) and expressing MHC I presenting the OVA peptide SIINFEKL There was a significant increase seen in K_2_ + 4.8 mM 1V209-K_2_ over all other groups.

Subsequently, we sought to investigate IgG subclasses to better characterize the effects of 1V209-K_2_. Although there are four IgG subclasses in both mice and humans, there are not clear inter-species analogs in subclass structure or function.^[46]^ A significant difference between mouse and human antibody production involves class switching, where human B cells are understood to switch sequentially from IgM to IgG3 to IgG1 to IgG2 to IgG4, while the prevailing theory in mice suggests a quartet model in which IgM can switch to IgG1, IgG2a, IgG2b, or IgG3.^[47]^ In this model, all four murine IgG exist together and function in concert, despite having different and sometimes opposing effects. In this study, the four subclasses were measured by ELISA from serum collected at 12 weeks post-vaccination (Figure 7B). The two carrier-free groups showed OVA-specific titers below the limit of detection for all subclasses, correlating with negligible pan-IgG titers measured throughout the study. No significant differences in the levels of OVA-specific IgG3 were observed between any of the hydrogel groups while both 1V209-containing hydrogels had similar IgG1 levels to alum and significantly higher IgG1 than unmodified K_2_. However, there were significant increases in OVA-specific IgG2a and IgG2b titers in both 1V209-K_2_-containing groups compared to both K_2_ and alum. Notably, both IgG2a and IgG2b exhibit strong Fc-gamma receptor (FcγR) binding and are considered proinflammatory.^[48]^ IgG2a (or its analog, IgG2c, in C57BL/6 mice) and IgG2b have been previously demonstrated to be upregulated in the presence of TLR7 signaling,^[49–51]^ providing a potential mechanism for this response in 1V209-K_2_-adjuvanted formulations. While IgG2b is T-cell independent, the murine IgG2a response has been shown to be upregulated by interferon-γ and correlated to Th1-mediated immunity, fitting with the expected Th1 response induced by TLR7 agonists like 1V209. The IgG1:IgG2a ratio is also significantly lower in the two 1V209-containing MDP groups than in alum; both K_2_ + 4.8 mM 1V209-K_2_ and K_2_ + 0.1 mM 1V209-K_2_ have a lower average ratio than K_2_, but only the difference between K_2_ + 4.8 mM 1V209-K_2_ and K_2_ is statistically significant (Figure 7C). As IgG1 and IgG2a correlate to Th2 and Th1 immunity, respectively, an IgG1:IgG2a ratio near unity is thought to indicate a balanced Th2/Th1 immune response.^[52]^

To further investigate the immune response, mice were euthanized one week after vaccination and their splenocytes were analyzed by flow cytometry. OVA delivered in K_2_ + 4.8 mM 1V209-K_2_ hydrogels induced a significant increase in CD3^-^CD86^+^ cells compared to alum, K_2_, and 0.1 mM 1V209, although K_2_ + 0.1 mM 1V209-K_2_ and OVA alone were also slightly elevated (Figure 7D); CD86 is a costimulatory molecule on APCs that binds to CD28 on T cells, which induces proliferation and differentiation.^[53]^ These data may be correlated with a statistically insignificant decrease in the CD4:CD8 ratio (Figure 7E) and a slight increase in CD69-activated CD8^+^ T cells (Figure 7F).

Measuring the specificity of APCs for OVA using commercially available reagents requires the expression of the H-2K^b^ MHC I haplotype, which is expressed by C57BL/6 mice but not BALB/c mice.^[54]^ Because measuring this population would be very valuable in this (and other) studies, we sought to use it in our SKH1-Elite mice, whose MHC I haplotype had not previously been determined. Using a haplotype-specific anti-H-2K^b^ antibody, we observed that all SKH1-Elite mice tested expressed H-2K^b^, using C57BL/6 and BALB/c mice as positive and negative controls, respectively (Figure S12). Given this confirmation, we measured antibody binding to MHC I H-2K^b^ bound to SIINFEKL, an epitope of OVA. We observed a significant increase in CD45^+^CD3^-^ cells expressing this marker in mice receiving K_2_ + 4.8 mM 1V209-K_2_ compared to all other groups (Figure 7G), suggesting that a high concentration of 1V209 induces non-T cell leukocytes to take up and present antigen to cytotoxic T cells in the spleen.

Tissue sections containing hydrogels excised 1 and 13 weeks after vaccination were immunohistochemically stained for F4/80^+^ macrophages, CD11c^+^ dendritic cells, and TNF-α, a pro-inflammatory cytokine. At the week 1 timepoint, some dendritic cells were visible in the inflamed skin surrounding the 1V209-containing hydrogels, while a noticeable ring of DCs was observed infiltrating the K_2_ gel (Figure 8). While we observed a relatively small number of macrophages at the edges of alum and no clear F4/80 staining in K_2_, there was clear infiltration of large numbers of macrophages into the skin surrounding both 1V209-modified hydrogels, with greater staining in the K_2_ + 4.8 mM 1V209-K_2_ group. We hypothesize that the greater concentration of macrophages in K_2_ + 4.8 mM 1V209-K_2_ hydrogels is correlated to the significant increase in cells with MHC I presenting SIINFEKL at 1 week seen in flow cytometry. Finally, the region of the 1V209-modified hydrogels with infiltrated cells appears to stain almost completely positive for TNF-α, with a smaller amount of staining in the group receiving K_2_ and no noticeable staining in the group receiving alum. TNF-α is secreted by Th1 immune cells and is involved in the initiation of adaptive immune responses. This finding suggests that the TLR7 agonist-conjugated MDPs are interacting with TLR7 molecules in innate immune cells to drive a Th1 adaptive immune response.

**Figure 8.**
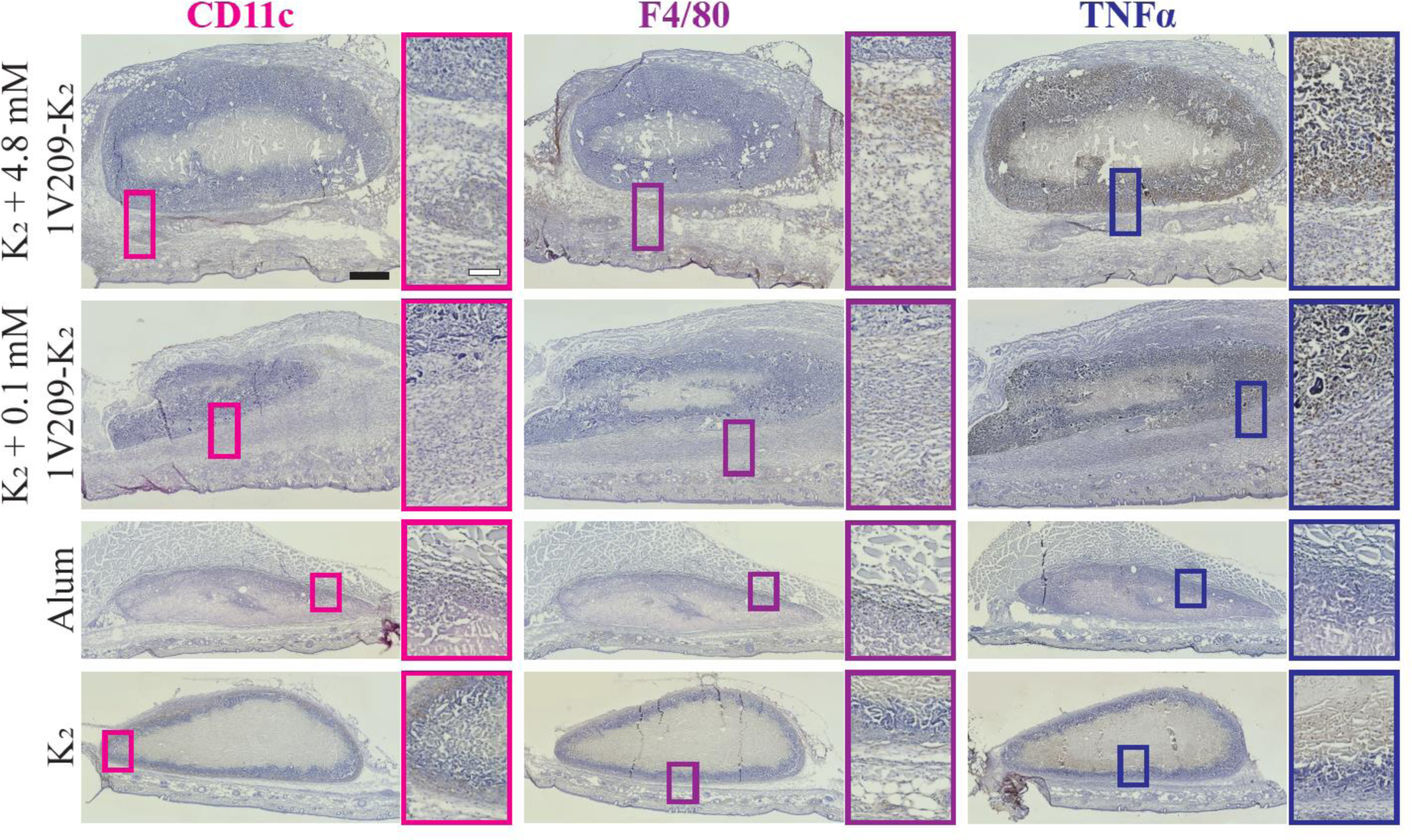
Immunohistochemical analysis of hydrogels excised one week after subcutaneous injection. Positive staining appears brown; nuclei are stained blue. Some infiltration of CD11c dendritic cells is visible in the skin around the 1V209-containing hydrogels, with slight infiltration into alum and a clear ring at the edge of K_2_. F4/80 macrophages are visible to a qualitatively greater degree in skin surrounding 1V209-containing hydrogels over alum and K_2_. Heavy TNF-α staining is visible within both 1V209-containing hydrogels, with slight staining in K_2_ and no visible staining in alum. The black scale bar signifies 500 µm for all full images, while the white scale bar denotes 100 µm for all insets.

Thirteen weeks after injection, the comparably small remaining K_2_ hydrogels had no noticeable staining of CD11c or F4/80 and displayed only diffuse TNF-α staining (Figure 9). The K_2_ + 0.1 mM 1V209-K_2_ group showed limited populations of dendritic cells and some diffuse macrophages, while we observed widespread infiltration of both dendritic cells and macrophages in the K_2_ + 4.8 mM 1V209-K_2_ group. TNF-α signal decreased considerably in 1V209-containing hydrogels between 1 and 13 weeks. We hypothesize that the enhanced or extended infiltration of dendritic cells and macrophages into the K_2_ + 4.8 mM 1V209-K_2_ hydrogels, relative to the K_2_ + 0.1 mM 1V209-K_2_ group, are correlated to the lower IgG1:IgG2a ratio and therefore stronger relative Th1 response seen at week 12.

**Figure 9.**
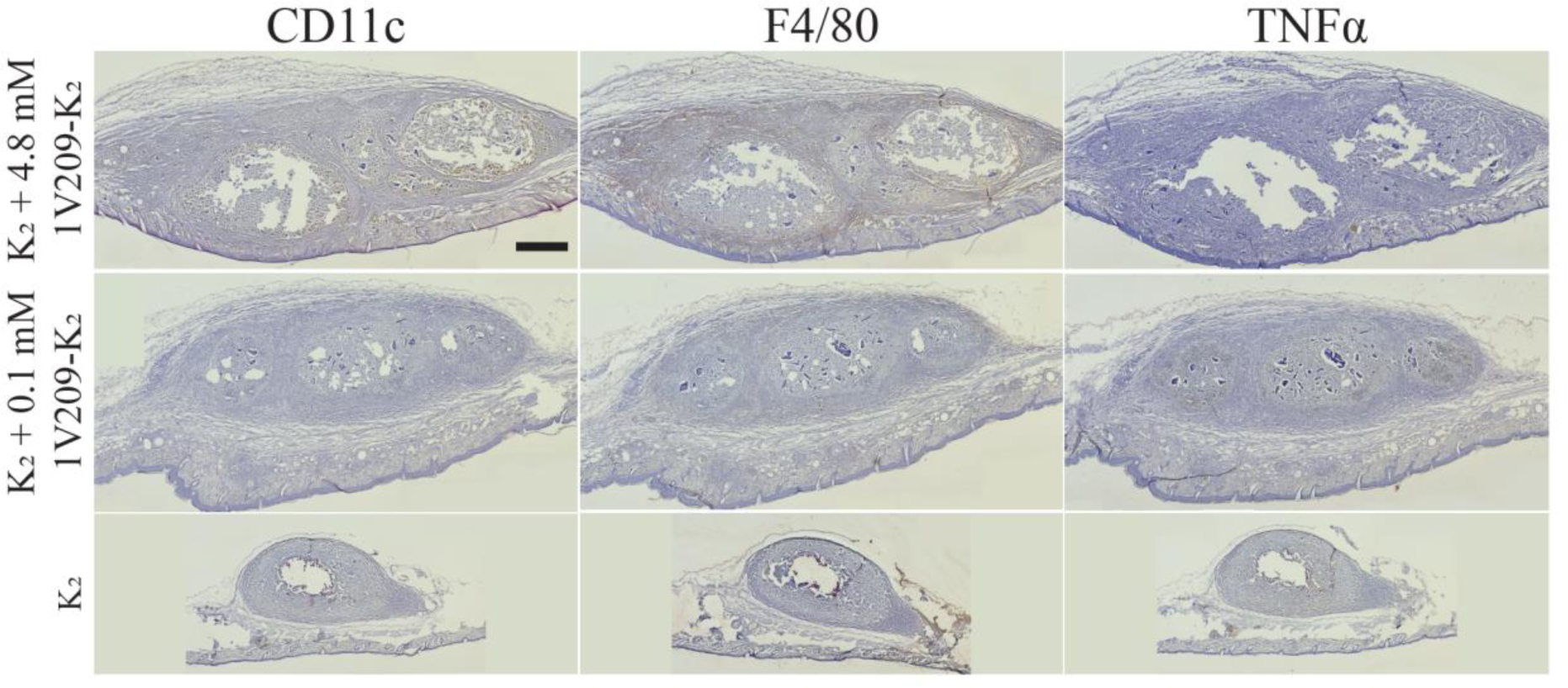
Immunohistochemical analysis of hydrogels excised 13 weeks after subcutaneous injection. Positive staining appears brown; nuclei are stained blue. There is clear infiltration of CD11c dendritic cells and F4/80 macrophages into K_2_ + 4.8 mM 1V209-K_2_ hydrogels, with minimal TNF-α staining compared to the one-week timepoint. K_2_ + 0.1 mM 1V209-K_2_ shows some F4/80 and TNF-α staining within the gel, but minimal CD11c. The comparatively small remaining K_2_ hydrogel also showed minimal staining in CD11c and F4/90 and some slight staining with TNFα. Alum is not pictured as the hydrogels were unable to be located within the subcutaneous space. The black scale bar signifies 500 µm.

## Conclusions

In this study, we show that the conjugation of 1V209 to K_2_ acts as an adjuvant, significantly improving the humoral immune response to a co-delivered model antigen in mice compared to a dose-matched injection of 1V209 or unmodified K_2_. We demonstrate that this improvement is likely a result of persistence of 1V209 with antigen at the site of injection, prompting an increased inflammatory response and recruitment of macrophages and dendritic cells. A clear enhancement of TNF-α production was observed in 1V209-containing gels compared to unmodified K_2_ one week after injection and an increase in TNF-α-producing cells has been correlated to stronger immunity in vaccinations against malaria^[55]^ and group A streptococcus.^[56]^ 1V209-modified K_2_ hydrogels induced significantly higher IgG2a and IgG2b titers than alum 12 weeks after immunization, in turn decreasing the IgG1:IgG2a ratio suggestive of a more balanced Th2/Th1 immune response. This is significant because K_2_ hydrogels without 1V209 modification were recently shown to strongly bias the immune response toward humoral immunity in C57BL/6J mice.^[37]^ This platform allows for modulation of the immune response toward protective Th1 immunity while retaining the benefits of MDPs, including extended release of antigen and shear-thinning properties for facile injection, making it an attractive adjuvant for infectious disease vaccines.

## Acknowledgements

The authors thank Alyssa Kunkel, Mei-Li Laracuente, and Sam Wu for their assistance. B.H.P. received funding from the National Science Foundation Graduate Student Research Fellowship program and the National Cancer Institute F99/K00 program (award number: F99CA284262). This work was supported by National Institute of Health grants R01 DE030140, K22AI146215, and R35GM143101, the Welch Foundation grant C-2141, the Cancer Prevention and Research Institute of Texas grant RR190056, and start-up funds from Rice University. The table of contents figure and Figure 1 were made using Biorender.com.

