## Supplemental Information for "A TLR7 Agonist Conjugated to a Nanofibrous Peptide Hydrogel as a Potent Vaccine Adjuvant"

| Stain | Company | Product No. | Volume for 1M Cells |
|---|---|---|---|
| Anti-CD45 -Pacific Blue | BioLegend | 103126 | 1 µL |
| Anti-CD3 -AlexaFluor 700 | BioLegend | 100216 | 2 µL |
| Anti-CD4 -AlexaFluor 488 | BioLegend | 100529 | 0.5 µL |
| Anti-CD8 -APC/Fire750 | BioLegend | 100766 | 1 µL |
| Anti-CD69 -Brilliant Violet 605 | BioLegend | 104530 | 1 µL |
| Anti-CD86 -Brilliant Violet 421 | BioLegend | 105031 | 1 µL |
| Anti-H-2K$^b$ bound to SIINFEKL -APC | BioLegend | 141606 | 2.5 µL |
| LIVE/DEAD Fixable Yellow Stain | Thermo Fisher | L34959 | 1 µL |

**Table S1.** Flow cytometry markers.

| Stain | Company | Product No. | Dilution |
|---|---|---|---|
| Anti-TNF-α –Biotin | BioLegend | 506311 | 1/100 |
| Anti-F4/80 –Biotin | BioLegend | 123105 | 1/100 |
| Anti-CD11c –Biotin | BioLegend | 117303 | 1/100 |
| HRP Streptavidin | BioLegend | 405210 | 1/50 |

**Table S2.** Immunohistochemistry markers.

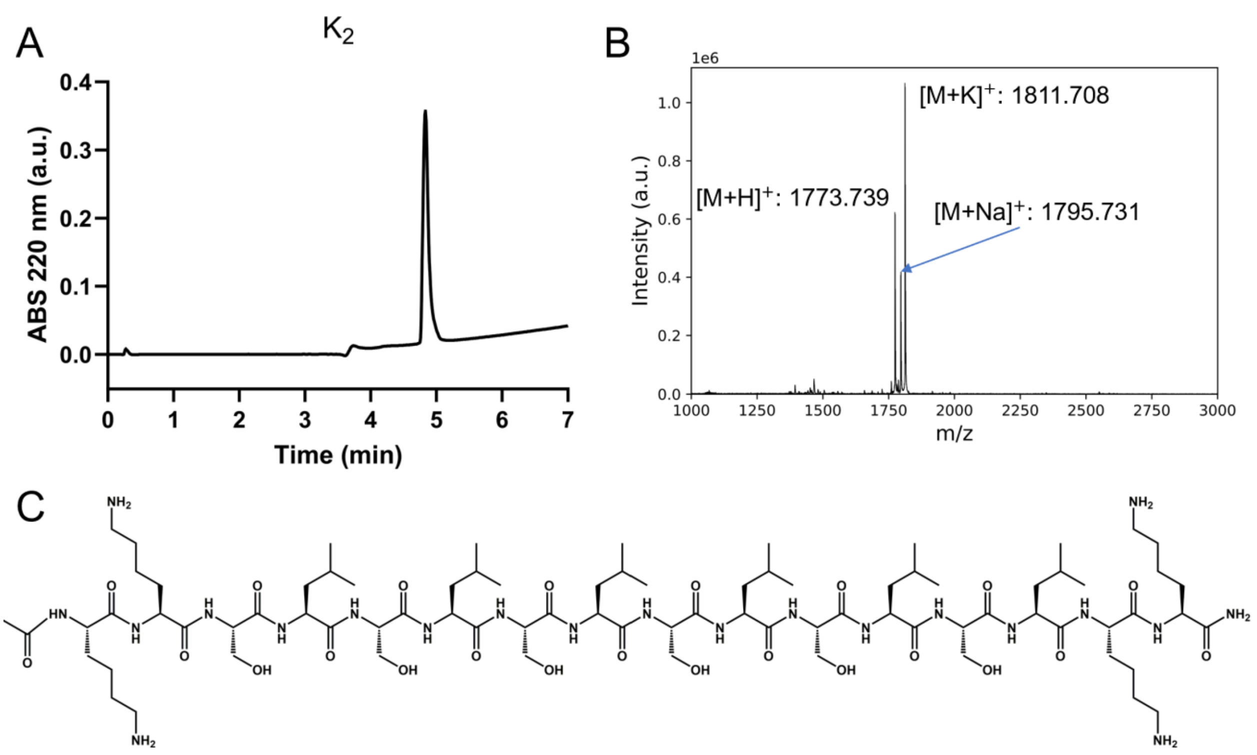


**Figure S1.** Characterization of the purified $K_2$ peptide by A) UPLC and B) MALDI-MS. C) The full chemical structure of $K_2$.

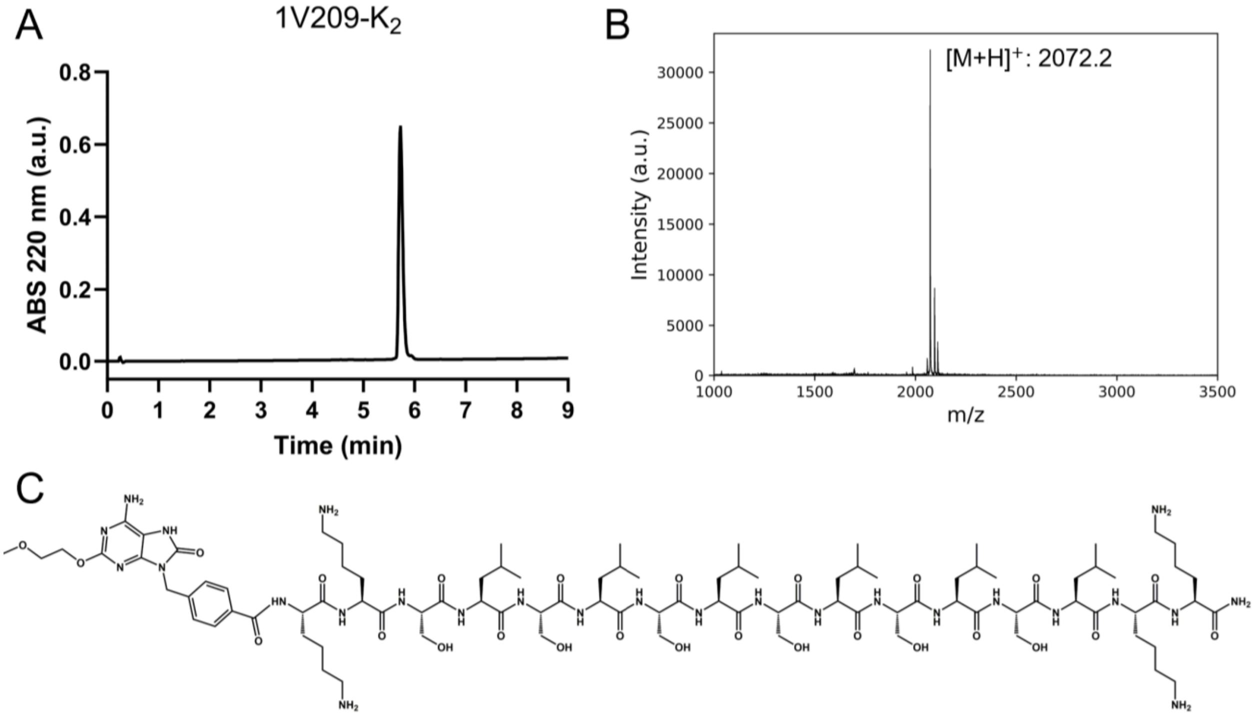


**Figure S2.** Characterization of the purified 1V209-$K_2$ peptide, also called MDP1, by A) UPLC and B) MALDI-MS. C) The full chemical structure of 1V209-$K_2$.

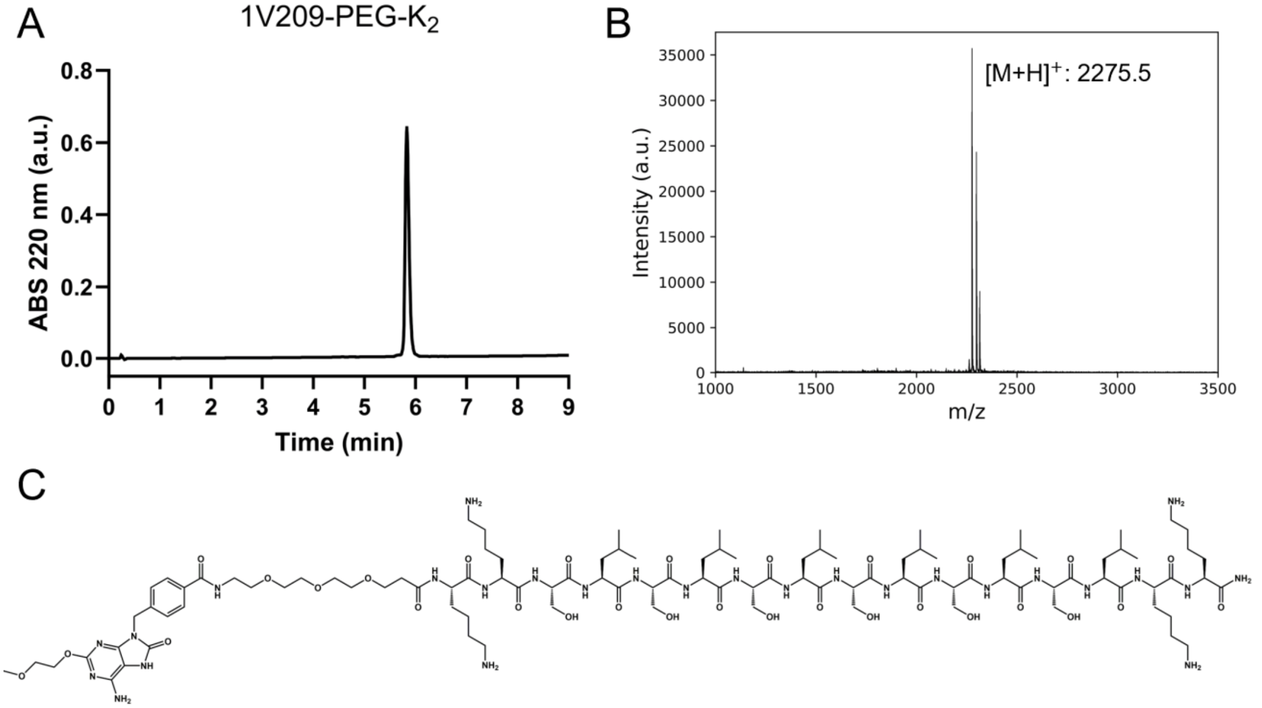


**Figure S3.** Characterization of the purified 1V209-PEG-$K_2$ peptide, also called MDP2, by A) UPLC and B) MALDI-MS. C) The full chemical structure of 1V209-PEG-$K_2$.

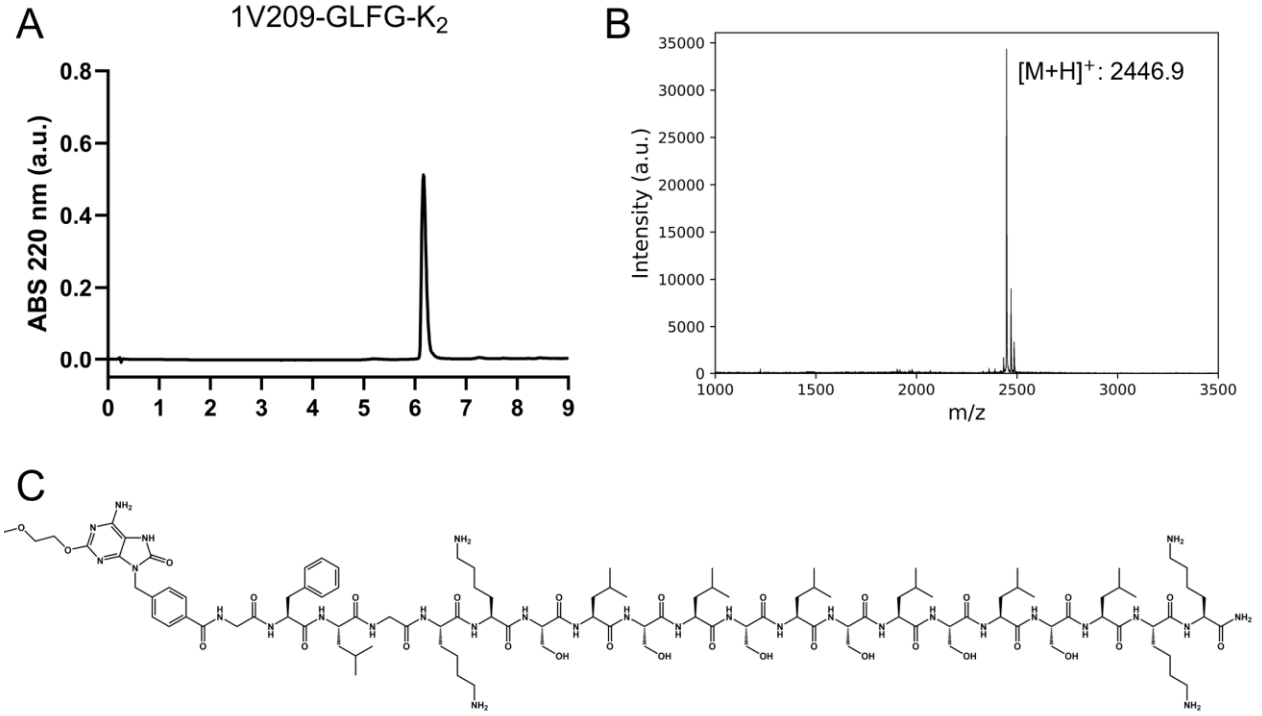


**Figure S4.** Characterization of the purified 1V209-GLFG-$K_2$ peptide, also called MDP3, by A) UPLC and B) MALDI-MS. C) The full chemical structure of 1V209-GLFG-$K_2$.

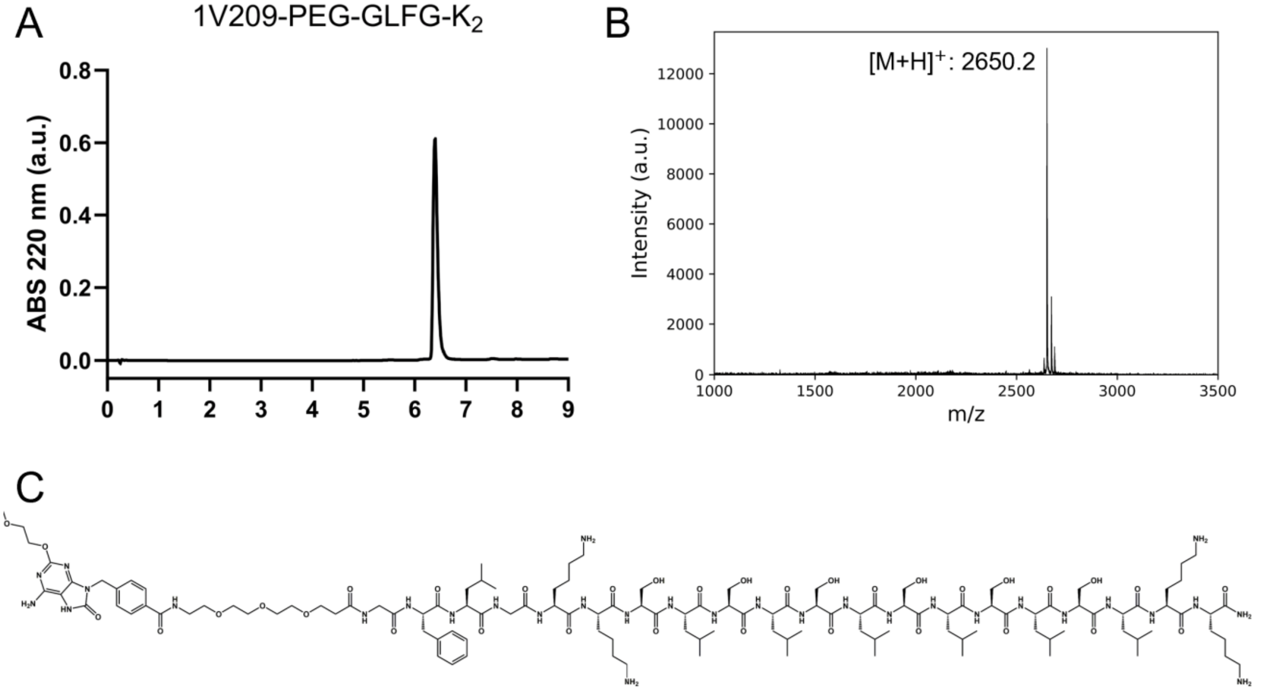


**Figure S5.** Characterization of the purified 1V209-PEG-GLFG-$K_2$ peptide, also called MDP4, by A) UPLC and B) MALDI-MS. C) The full chemical structure of 1V209-PEG-GLFG-$K_2$.

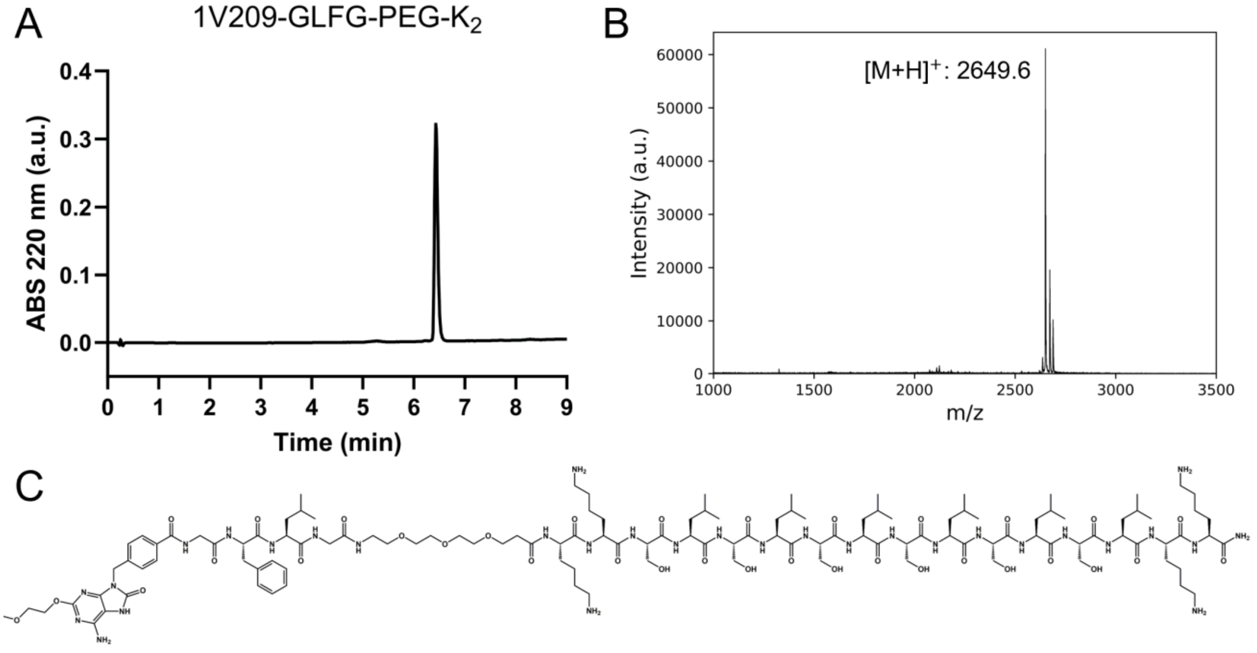


**Figure S6.** Characterization of the purified 1V209-GLFG-PEG-$K_2$ peptide, also called MDP5, by A) UPLC and B) MALDI-MS. C) The full chemical structure of 1V209-GLFG-PEG-$K_2$.

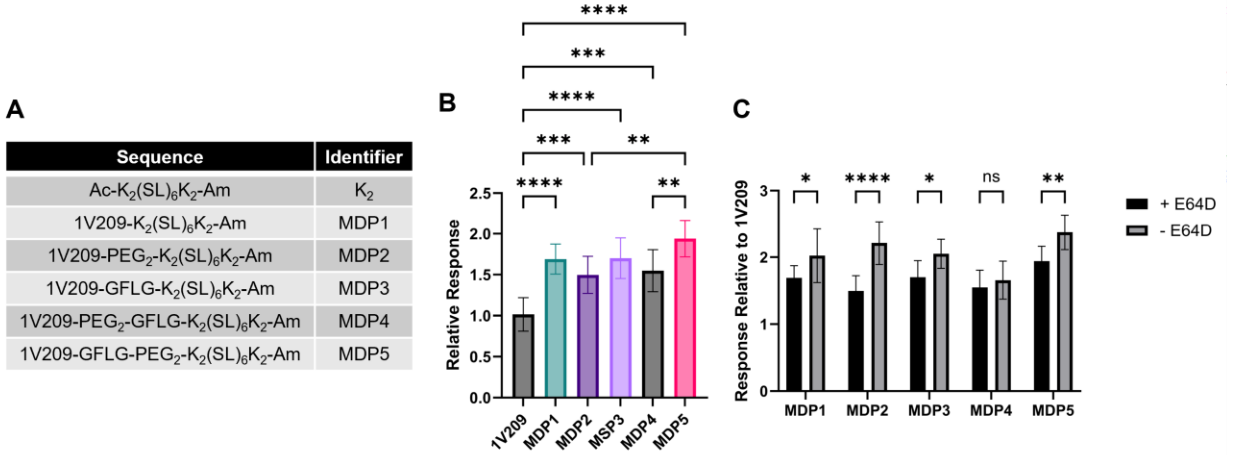


**Figure S7.** A) The amino acid sequences of each synthesized MDP. B) The relative TLR7 agonist activity of each MDP, normalized to the response of unconjugated 1V209 when treated with the cysteine protease inhibitor E64D. C) The TLR7 agonist activity of each MDP with and without the E64D inhibitor.

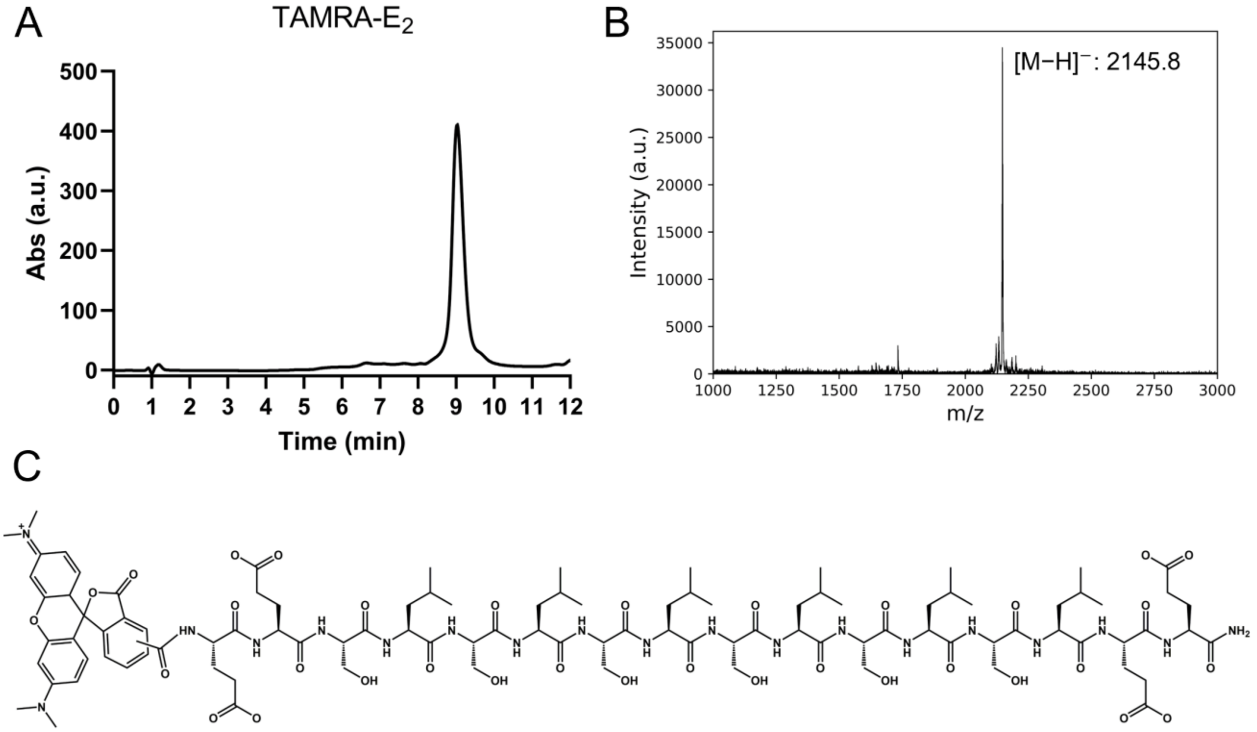


**Figure S8.** Characterization of the purified TAMRA-$E_2$ peptide by A) UPLC and B) MALDI-MS. C) The full chemical structure of TAMRA-$E_2$.

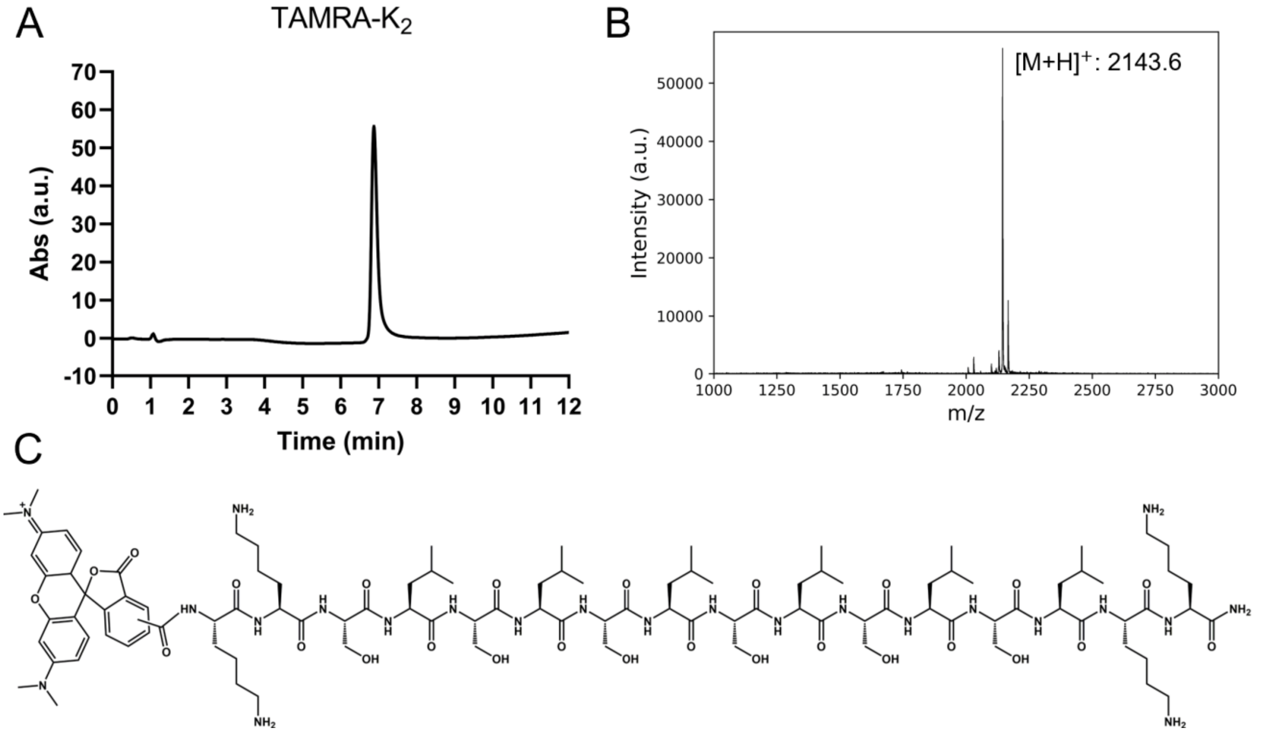


**Figure S9.** Characterization of the purified TAMRA-$K_2$ peptide by A) UPLC and B) MALDI-MS. C) The full chemical structure of TAMRA-$K_2$.

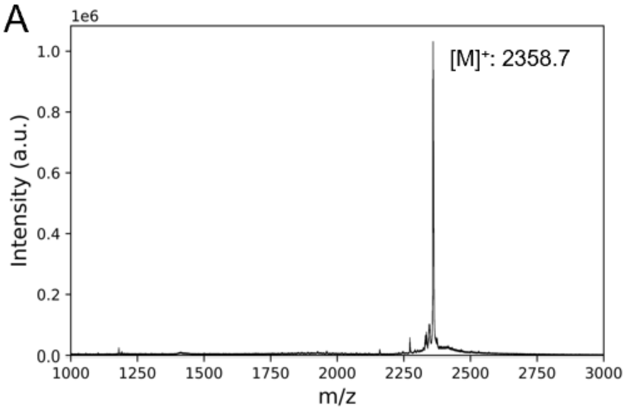


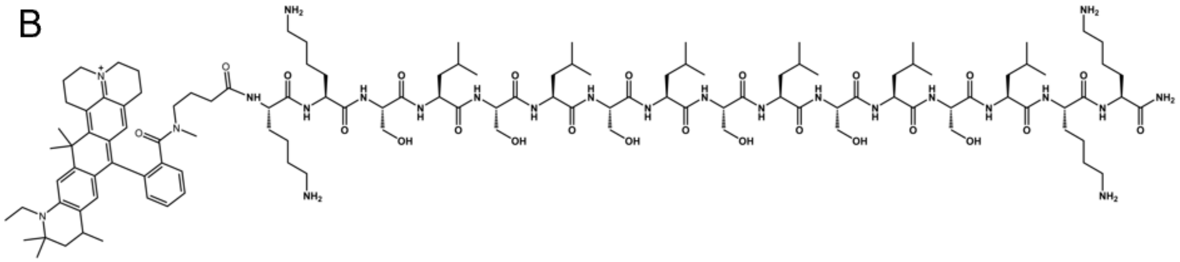


**Figure S10.** Characterization of the purified ATTO647N-$K_2$ peptide by A) MALDI-MS. B) The full chemical structure of ATTO647N-$K_2$.

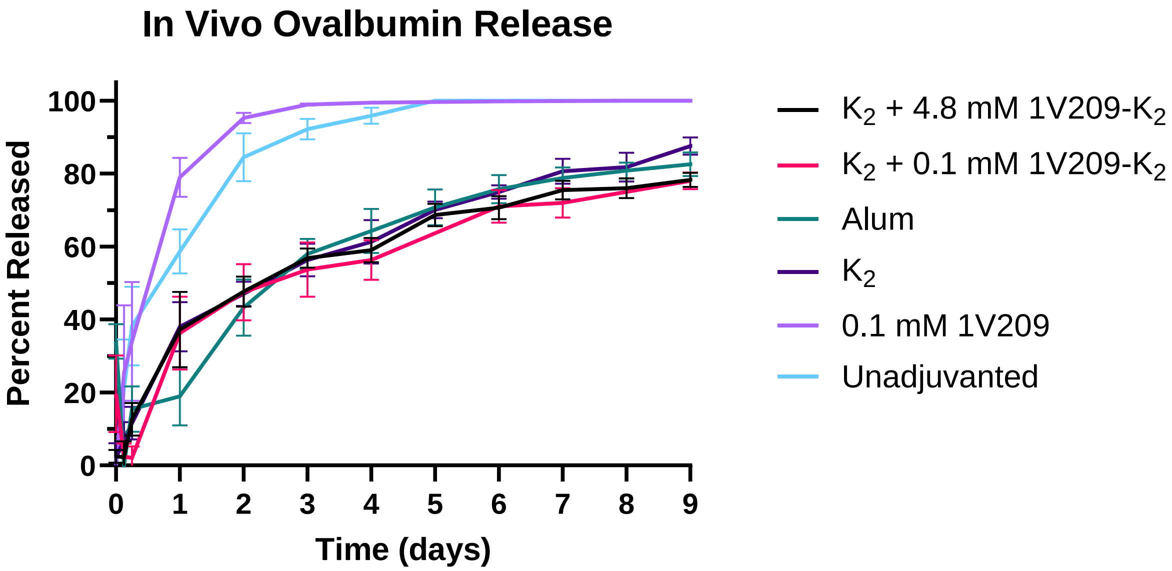


**Figure S11.** In vivo release of fluorescently labeled OVA from $K_2$ + 0.1 mM 1V209-$K_2$ and from 0.1 mM 1V209 over nine days, as compared to the remaining groups. OVA was released from $K_2$ + 0.1 mM 1V209-$K_2$ at a similar rate to $K_2$ + 0.1 mM 1V209-$K_2$. OVA cleared from the 0.1 mM 1V209 injection site slightly faster than from injection sites receiving OVA alone.

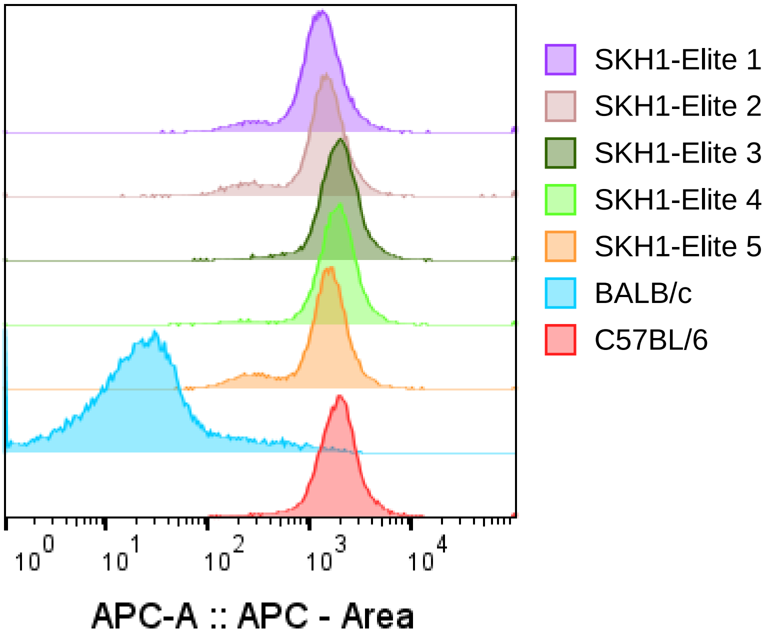


**Figure S12.** Flow cytometry measurements of anti-mouse MHC I H-$2K^b$ conjugated to the fluorophore APC. All five SKH1-Elite mice from five different litters showed similar levels of staining as the positive control C57BL/6 mouse, while the negative control BALB/c was greatly reduced.
